# Whole brain fluorescence imaging in *Drosophila* reveals spreading depression and its initiation, propagation, and resilience dynamics

**DOI:** 10.1101/2025.10.09.681455

**Authors:** A. L. Harris, G. Crowe, E. Nelson, M. Al Mokbil, W. Stein

## Abstract

Spreading depression (SD) is a wave of neuronal hyperactivity followed by depolarization block that propagates through large brain regions and is associated with disorders such as migraine, stroke, and brain injury. The mechanisms that initiate SD and alter susceptibility to it remain incompletely understood. Here, we use whole-brain fluorescence imaging with genetically encoded pan-neuronal calcium and voltage sensors to observe SD in *Drosophila melanogaster*. We show that rapid cooling, a naturally occurring environmental condition, as well as elevated extracellular potassium reliably elicit SD in both adult and larval flies. SD was characterized by a rapid and large rise in intracellular calcium that was accompanied by neuronal depolarization and stark changes in the transperineuronal potential. In adults, SD occurred at 6.7 ± 0.6°C (N=15) and in larvae at 6.0 ± 0.3°C (N=30). SD initiation was not restricted to specific sites, but initiated at multiple, variable sites across and within individuals, with an average of 3.0 ± 0.7 (N=8) initiation points per brain. In all cases, SD spread throughout large areas of the nervous system. In a high-throughput larval assays that allows the simultaneous monitoring of up to 16 animals with repeated cooling cycles, we demonstrate that single SD events are followed by a transient refractory period lasting up to 45 minutes, during which the threshold for subsequent SD was significantly elevated. This was the case in adult and larval brains of all developmental stages. The refractory effect was independent of neuronal depolarization, suggesting that homeostatic processes alter SD susceptibility following an initial SD event. Taken together, our findings demonstrate that SD initiation and propagation are not restricted to specific regions, neuronal populations, or developmental stages, and they reveal fundamental properties of adaptive changes to SD susceptibility in a genetically tractable model. Building upon the extensive genetic toolkit available in *Drosophila*, this work establishes the fly as a complementary model for understanding conserved cellular and circuit-level mechanisms of SD relevant to human neurological disorders.

## I. Introduction

Spreading depression (SD) is a dramatic neurophysiological phenomenon that occurs during disorders such as migraine, stroke, cerebral ischemia, and brain injury^1^. During SD, a wave of neuronal hyperexcitability and subsequent inactivation propagates through large areas of the brain (often cortex) at a speed of a few millimeters per minute^2^. At a cellular level, SD is characterized by rapid neuronal firing followed by a loss of spiking due to depolarization block. It typically manifests with negative symptoms, but has recently been hypothesized to have a protective effect against more severe disorders^3^. Studies in mammalian brains, alongside computational models, have successfully characterized the dynamics and physiological mechanisms of SD spread^1,2,4–8^. These include dramatic changes in intra- and extracellular ion concentrations, such as elevated extracellular potassium and intracellular sodium. The elevated potassium concentration, in particular, propagates at a characteristic slow velocity through the extracellular medium via diffusion, mediating the spread of the hyperactivity and subsequent depolarization block. The consequences of this disrupted ion homeostasis include cell swelling, altered blood flow, inflammation, and in the case of migraine, headaches (for review, see^1^).

Despite decades of study on cellular-level dynamics and SD spread^1,2,4–8^, the mechanisms that initiate, modulate, and affect SD susceptibility remain elusive. Current mammalian systems for studying SD struggle to provide large-scale and high-throughput recordings from intact brains that can identify the underlying mechanisms that alter SD susceptibility. Most commonly, studies of SD have imaged small brain regions with *in-vivo* experiments (e.g. ^9,10^) or elicited SD in *in-vitro* brain slices using artificial stimuli (e.g. ^11–13^), such as high focal potassium application or electrical or mechanical brain stimulation.

In contrast, insect nervous systems offer a more tractable system in which to study SD in fully intact animals. Recently, chill coma, a reversible state of paralysis that occurs in many chill-susceptible insects when exposed to low temperatures, has been proposed to be associated with SD^14–19^. This naturally occurring, experimentally controllable system enables the study of SD susceptibility through a process that is part of the animal’s natural experience. Initial work linking SD with chill coma has shown a dramatic extracellular potassium accumulation as insects are cooled to a few degrees Celsius^17–19^, accompanied by neuronal hyperactivity. Additionally, measurements of the DC field potential at two separate sites show a sudden drop^14,17,18^ that occurs with some time delay and has been interpreted as a spreading wave. However, no direct evidence of large-scale spread has yet been observed or measured, and it remains unclear whether the changes in field potential are correlated or occur independently. These studies suggest a link between SD and chill coma, and show that locally, the ion concentration and electrophysiological changes observed in insects are similar to those in mammals^20^. However, electrophysiological and ion concentration measurements are technically challenging, low-throughput, and provide only a limited picture of the physiological dynamics of SD initiation, spread, and susceptibility.

We introduce a novel method for observing and quantifying SD in *Drosophila melanogaster* using whole brain fluorescent imaging in which both the spatial and temporal dynamics of SD can be measured. We present evidence that SD can be elicited using rapid cooling and extracellular potassium application, and show that SD spreads across large areas of the nervous system. Our technique allows us to capitalize on the many advantages of the *Drosophila* study system, including targeted gene expression and high-throughput imaging techniques^21–26^. Using genetically encoded pan-neuronal calcium and voltage sensors, we show that SD is not restricted to particular brain areas but is initiated at different and multiple sites that vary across and within animals. We also show that SD exists not only in adult *Drosophila*, but also in all three larval stages, providing further evidence for the ubiquity of SD across widely varying neuronal connectivities. Our assays for SD measurements in larvae provide a simple and high-throughput technique for observing SD in up to 16 animals simultaneously, enabling the rapid testing of SD initiation stimuli. Through repeated exposure to cold in our high-throughput assay, we demonstrate that a singular SD event confers protection for up to 45 minutes against future SD events, during which time the threshold to elicit SD is elevated. This refractory period is reminiscent of a similar feature observed in mammals^27,28^, but is shown here for the first time in fully intact animals under naturally experienced conditions. Overall, our results demonstrate that SD exists and spreads across large brain regions in adult and larval *Drosophila*, indicating that SD initiation and propagation are not restricted to specific regions, neuronal populations, or developmental stages. They also show that single SD events lower the susceptibility to future SD events, confering resilience for up to 45 minutes, and establishes the fly as a complementary model for understanding dissect conserved cellular and circuit-level mechanisms of SD.

## II. Results

### Cooling elicits a slowly spreading wave of neuronal hyperexcitation in adult fly brains

Spreading depression is associated with an initial high neuronal activity that is followed by a long-lasting phase of activity depression due to the continuous depolarization of the neurons and a subsequent inactivation of the sodium channels that mediate action potentials. Once triggered in a localized region, this depression then spreads across large areas of the nervous system through an effect thought to be associated with, and mediated by, large changes in extracellular potassium concentrations^2^. We tested the hypothesis that spreading depression exists in adult fly brains and can be elicited by rapid cooling. Previous studies^15,17^ have indicated that this is the case but relied on low-throughput electrophysiology with only indirect indications of spread and no ability to measure or observe the dynamics of the spreading wave of depolarization. To definitively determine whether rapid cooling of the fly brain initiates spreading depression, we pan-neuronally expressed the calcium indicator GCaMP6m (see Materials and Methods) and measured its fluorescence while rapidly cooling the adult fly brain. The fluorescence of genetically encoded calcium indicators, such as GCaMP6m, correlates with neuronal depolarization and is particularly reliable in indicating the rise of neuronal activity^29^.

We mounted adult flies (aged 17 to 67 days) and exposed the brain by microdissection of the posterior head capsule cuticle (Figs. 1A,B). This allowed us to record large-scale neuronal activity as evidenced by spontaneously occurring changes in fluorescence in the different brain regions (Figs. 1C,D). For example, in Fig. 1D, we detected spontaneous fluorescence changes at room temperature at six different locations across the mushroom body. The fluorescence traces of the left mushroom body showed correlated, but propagating activity that was independent of that observed in the right mushroom body. Overall, the changes in detected spontaneous GCaMP6m fluorescence values ranged between 3 and 50 % of the background fluorescence of the location where they were measured (see also supplemental 1). Brains in which no spontaneous activity was detected were considered damaged during dissection and discarded. Brains with spontaneous activity were cooled from room temperature (20 - 22°C) to near 0°C. We tested cooling rates between 0.8°C/min and 3°C/min, but for each experiment, the rate of cooling was kept constant. In all cases, cooling was followed by a warming back to room temperature at a rate of 3°C/min. Figure 2A shows a representative example of the observed fluorescence during this protocol. Panel i shows the fluorescence after the brain had been cooled to 13°C. Panel ii shows the same brain after further cooling to 5.9°C, and that a slight overall increase in fluorescence was observed throughout the brain. Panel iii shows that there was a large increase in fluorescence at 2°C that was particularly prevalent in the mushroom body. This occurred just before the coldest temperature of 1.4°C. Panel iv (4.6°C) shows that the fluorescence decreased slowly during the initial re-warming phase. Finally, panel v shows that the fluorescence further decreased when the brain was warmed back to 18°C.

**Figure 1.**
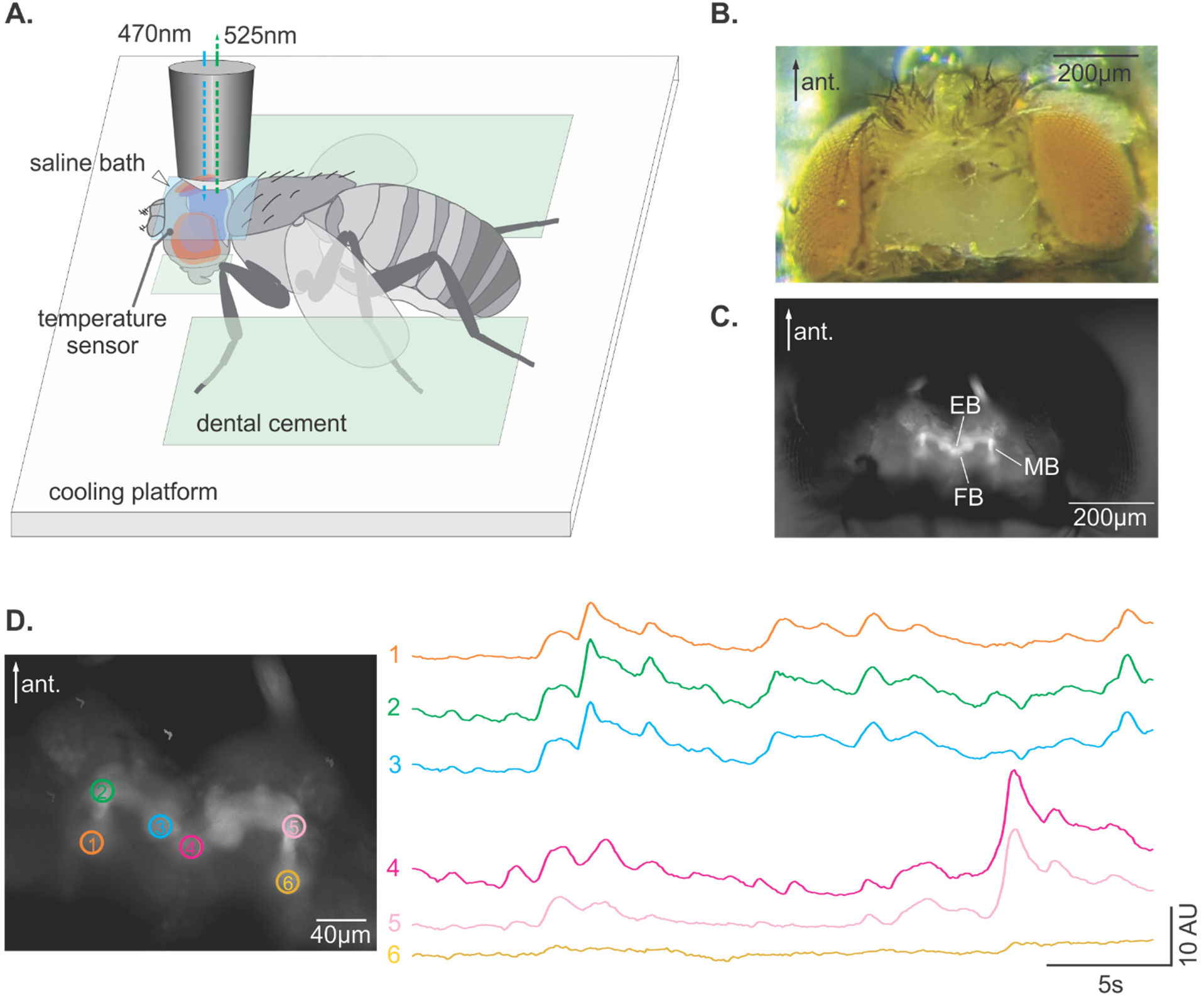
Whole brain fluorescent imaging of adult flies. A) Schematic of experimental setup to rapidly cool from room temperature to near 0℃ while measuring GCaMP6m or Arclight fluorescence in adult flies. B) Brightfield image of an adult fly head with the posterior head cuticle removed to expose the brain. C) Same fly as in (B), expressing pan-neuronal GCaMP6m under fluorescent light (470 nm illumination, 525 nm detection). The mushroom body (MB), ellipsoid body (EB), and fan-shaped body (FB) are clearly visible. D) Left – regions of interest (ROIs) used to measure spontaneous activity in the brain at room temperature. Right – fluorescence (arbitrary units) measured in each ROI at room temperature. Traces are separated vertically for clarity. See also supplemental 1.

**Figure 2.**
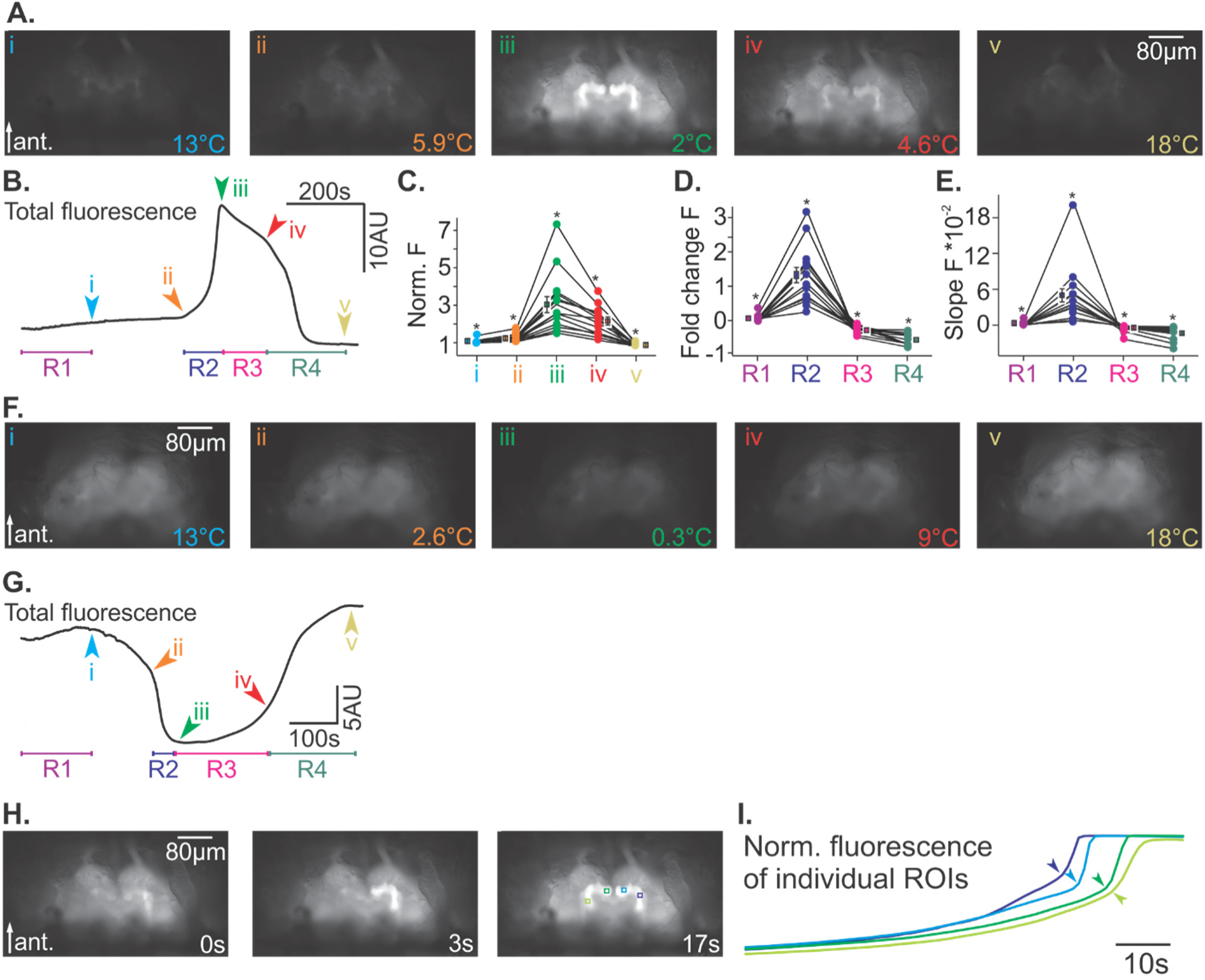
Changes in fluorescence during rapid cooling and rewarming can be classified into 4 distinct regimes. A) Representative original recording of GCaMP6m fluorescence during cooling from room temperature to near 0℃ and then rewarming back to room temperature. i) End of regime 1, 13℃. ii) Start of regime 2 at 5.9℃. iii) End of regime 2 = start of regime 3 at 2℃. iv) End of regime 3 = start of regime 4 at 4.6℃. v) End of regime 4, at 18℃. B) Representative whole brain GCaMP6m fluorescence over time showing the 4 distinct regimes (R1 – R4) and their starting and ending points (i – v). C) Quantification of normalized GCaMP6m fluorescence at points i – v. For each animal, the mean fluorescence was normalized to the mean fluorescence value at 18 °C at the start of the cooling ramp. The normalized fluorescence value at each point was significantly different from each of the other points (Friedman chi-square test, N = 15, χ² = 52.000, dF = 4, P<0.001, Student-Newman-Keuls posthoc test at P<0.05). D) Fold change in GCaMP6m fluorescence during each of the regimes. The change in fluorescence in each regime is significantly different from each of the other regimes (Friedman chi-square test, N = 15, χ² = 39.000, dF = 3, P<0.001, Student-Newman-Keuls posthoc test at P<0.05). E) Slope of GCaMP6m fluorescence change over time during each of the regimes. The slope of fluorescence change during each regime was significantly different from each of the others (Friedman chi-square test, N = 15, χ² = 39.000, dF = 3, P<0.001, Student-Newman-Keuls posthoc test at P<0.05). F) Same as (A), but for Arclight fluorescence. Temperatures at each point are indicated in the figure. G) Same as (B), but for Arclight fluorescence. H) Original recordings of GCaMP6m fluorescence during regime 2 showing that the fluorescence wave spreads through the mushroom body. Times of the frames are shown from the start of the spreading wave (t = 0). The right image shows 4 regions of interest (ROIs) used to measure the fluorescence in (I). See also supplemental 2. I) Normalized fluorescence for the ROIs shown in (H). The arrows indicate the start of regime 2 and the start of the SD event for each ROI. They are clearly separated in time, indicating that the fluorescence wave shows a spatial spread.

Figure 2B shows the mean fluorescence of the whole brain over the course of the experiment, starting at 18°C, cooling to 1.4°C, and then warming back to 18°C with the arrows and numbers indicating the panels shown in Fig. 2A. Figure 2C shows the quantification of the normalized fluorescence at each of the time points (i-v) for all animals (N = 15), demonstrating that the shape of the fluorescence curve in Fig. 2B was common across animals. In Fig. 2C, the mean fluorescence of each animal was normalized to its mean fluorescence value at 18°C at the start of the cooling ramp. A repeated measures Anova on ranks indicated a significant effect of time, and therefore temperature, on fluorescence with each point being significantly different from one another (Friedman chi-square test, N = 15, χ² = 52.000, dF = 4, P<0.001, Student-Newman-Keuls posthoc test at P<0.05). Notably, point iii showed the largest change, and on average reached 3.0 ± 0.4 times the fluorescence at 18°C.

In general, the observed fluorescence changes could be classified into 4 distinct regimes, which as we demonstrate below, correlate with expected physiological changes during a spreading depression event. In the first regime (R1, Fig. 2B, see also supplemental 2), as the temperature dropped from 18°C to 13°C, there was a slow and monotonic rise in fluorescence in all visible areas of the brain. This was indicative of increased depolarization across the brain. In regime 2, (R2, Fig. 2B), the fluorescence increased dramatically over much shorter time scales, beginning on average at 6.7 ± 0.6°C and peaking at 5.6 ± 0.6°C (N = 15), and taking 44.2 ± 5.6 seconds. This increase indicated a dramatic change in the state of the neurons, such as would be expected during rapid hyperexcitation. During regime 3, fluorescence slightly decreased at a slow rate (R3, Fig. 2B) even when cooling continued to an average coldest value of 3.7 ± 0.6℃ (N = 15). Finally, during regime 4, the fluorescence rapidly decreased to below its initial level as the temperature was increased back to 18°C (R4, Fig. 2B), suggesting that neuronal activity recovered from its hyperexcited state.

The fluorescence changes during the four regimes were all significantly different from one another, as shown by the fold change in fluorescence in Fig. 2D and the change in fluorescence over time (slope) shown in Fig. 2E. The most dramatic and rapid changes were again observed during regime 2, which showed an average 1.3 ± 0.2 fold increase from the beginning to the end of the regime (N=15, Fig. 2D) and a slope more than 50 times greater than during regime 1 (Fig. 2E).

The fluorescence signal thus strikingly resembled the expected shape of membrane polarization that neurons experience during spreading depression, i.e., a slow increase in voltage toward hyperexcitation followed by continuous depolarization with depressed activity, and ultimately a sudden recovery^2^. To test whether the observed fluorescence changes were indeed dependent on neuronal activity and not the result of an intrinsic temperature response of the GCaMP6m protein itself, we measured GCaMP6m fluorescence in homogenized brains. For this, we dissected out brains from 16 adult flies expressing GCaMP6m and homogenized them in 50 µL of calcium-free buffer solution (see Materials and Methods) to release the expressed GCaMP6m protein. 1 µL of the homogenized brain solution was added to 1 µL of 100 µM calcium buffer solution, and the changes in fluorescence of this brain solution were measured during cooling from room temperature to 0°C. We found that in contrast to intact brains, the GCaMP6m fluorescence of homogenized brains slowly and steadily decreased with colder temperatures (supplemental 3). Interestingly, fluorescence continued to decrease as the temperature was warmed back to room temperature, which might explain why in most experiments in intact brains the fluorescence at the end of the trial was slightly lower than at the beginning of the trial (e.g., Fig. 2B). However, there were no rapid changes in fluorescence of the homogenized brain solution that resembled regimes 2 and 3 of the intact brain. The largest fluorescence changes in intact brains were thus likely due to neuronal activity and not the result of an intrinsic temperature dependence of the GCaMP6m protein.

To further support that the observed GCaMP6m fluorescence changes in intact brains were due to neuronal activity, we repeated the intact brain experiments using a pan-neuronal expression of the voltage sensor Arclight (see Materials and Methods). Unlike GCaMP6m, which is only an indirect reporter of neuronal activity, Arclight fluorescence is dependent on the neuronal membrane potential. Specifically, Arclight fluorescence decreases with membrane depolarization^30^. Accordingly, when we repeated the cooling experiments, we observed a rapid drop in fluorescence during regime 2, indicative of a rapid depolarization of large numbers of neurons. Fig. 2F shows a representative example of the observed Arclight fluorescence during the cooling and rewarming protocol at the same milestones as shown in Figs. 2A. Overall, the Arclight fluorescence response was inverted from that of GCaMP6m. Panel i shows the fluorescence at 13°C. Further cooling caused a small drop in fluorescence shown in panel ii (2.6°C), followed by a larger and quicker decrease in fluorescence in panel iii (0.3°C). Upon warming, the fluorescence again increased (panel iv; 9°C) and continued to increase as the temperature returned to 18°C (panel v). Fig. 2G shows the mean fluorescence of the whole brain obtained during this trial across the four regimes defined in Fig. 2B. During regime 1, there was a slow increase in fluorescence. During regime 2, the fluorescence sharply dropped to its smallest value, indicative of a large-scale neuronal hyperexcitation. This drop occurred on average at 4.5 ± 0.4°C (N=7) and reached its minimum at 2.5 ± 0.3°C (N=7). The drop in fluorescence in regime 2 was followed by a slow increase in fluorescence during regime 3 as the temperature began to warm. Finally, in regime 4, there was a quick increase in fluorescence as the temperature returned to 18°C, suggesting an end to the hyperexcitation. Taken together, our Arclight and GCaMP6m data suggest that fly neurons undergo physiological changes resembling those observed during SD, with a sudden, long-lasting, and reversible depolarization that is triggered by cold temperatures.

Lastly, if the observed fluorescence changes in intact brains are caused by neuronal events during spreading depression, then they should propagate slowly across the brain. To determine if this was the case, we more closely inspected the GCaMP6m fluorescence dynamics of regime 2 (Figs. 2H,I) and found that the rapid rise in fluorescence did not occur simultaneously in all areas of the brain. Instead, there was a clear initiation point where fluorescence increased rapidly and then spread to neighboring areas. This wave moved slowly across large areas of the brain, leading to the increased mean brightness of the whole brain observed in regime 2. Figure 2H shows an example of a wave initiated in the right α lobe of the mushroom body that spread medially towards the β lobe, then jumped to the β lobe of the left mushroom body before ultimately reaching the left α lobe.

From our observations of fluorescence waves, we also noted that waves could be initiated in multiple locations. Fig. 3A shows an example where the first wave was initiated in the mushroom body of the right hemisphere, and a second wave was initiated shortly after in the mushroom body of the left hemisphere. These spreads occurred independently and were not contiguous. To detect initiation points of fluorescence spread, we placed a set of regions of interest (ROIs) along the areas of the brain that were affected by the spreading wave of fluorescence (Fig. 3Ai). We then plotted the normalized fluorescence intensity for each ROI over time (Fig. 3B) as a heatmap and detected the time point of maximum fluorescence (white trace in Fig. 3B). In the given example, the maximum fluorescence was reached at two sites (arrows) before it spread to neighboring ROIs. These two sites were thus classified initiation points. On average, we identified 3.0 ± 0.7 initiation points in each brain, with some initiation points leading to multiple spreads in different directions. On average, spreading waves traveled a distance of 63.7 ± 9.1 µm at a velocity of 12.8 ± 1.8 µm/s (N=40 spreads, 8 animals). Our measurements are likely to underrepresent the number of initiation points, because we did not monitor all brain areas (e.g., the optic neuropils were outside of our field of view, and we only measured a single focal plane). In some cases, we could see a rapid rise of the fluorescence in deeper areas of the brain before waves were visible or elicited in the monitored focal plane. In these cases, we could not identify how many additional separate initiation points were hidden from us in other focal planes.

**Figure 3.**
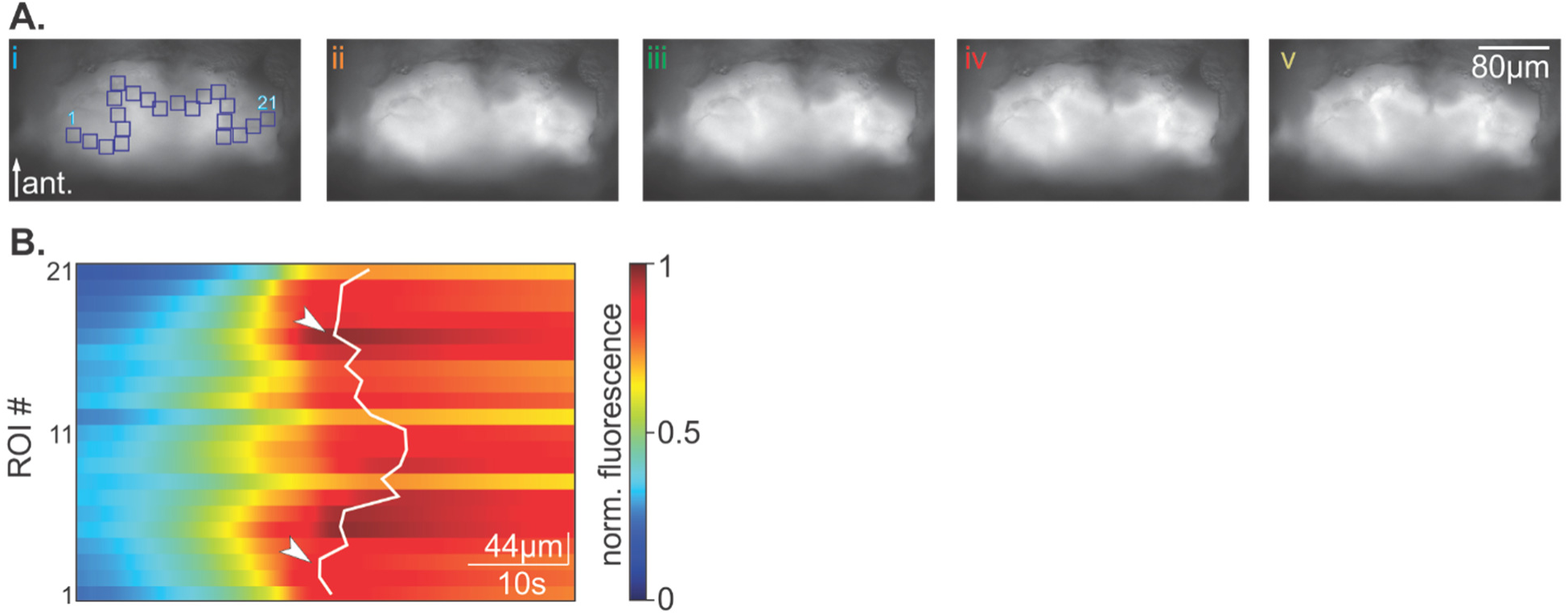
Multiple waves of spreading fluorescence were observed that started in different locations when the brain was cooled. A) Representative original recordings of GCaMP6m fluorescence showing that spreading waves were initiated in two different locations. i) Before spreading waves were initiated. 21 ROIs (blue squares) were drawn along the path that the spreading waves travelled. ii) Begin of spread in the mushroom body of the right hemisphere. iii) The high fluorescence spreads medially in the mushroom body of the right hemisphere. iv) A second wave of high fluorescence begins in the mushroom body of the left hemisphere while the first wave continues spreading medially in the right hemisphere mushroom body. v) The high fluorescence in the left hemisphere mushroom body spreads medially and laterally. B) Normalized fluorescence of each ROI defined in panel Ai over time. The white line traces the maximum fluorescence value of each ROI. The two points of wave initiation are indicated by the arrows.

Another hallmark of SD is that it is associated with a large increase in extracellular potassium that mirrors the hyperexcitation of the neurons and by itself causes further neuronal depolarization (following Nernst’s equation). Previous studies have shown that rapid cooling of insect brains can cause such a potassium increase and that this is reflected in changes of the transperineuronal potential^14,18^ (TPP). To determine whether the front of the observed fluorescence wave was associated with the expected changes in extracellular potassium concentration, we impaled the brain with a sharp microelectrode and recorded the TPP. Figure 4 shows the result of an experiment where the electrode was placed into the left brain hemisphere and the brain was subsequently cooled until a spreading fluorescence wave was observed. We found that there was a rapid change in TPP that coincided with the arrival of the wavefront of the GCaMP6m fluorescence (Fig. 4) at the recording site. This suggests that rapid changes in extracellular ion concentrations occurred at times when the neurons underwent strong cooling-induced depolarization, further supporting the hypothesis that cold temperatures elicit SD.

**Figure 4.**
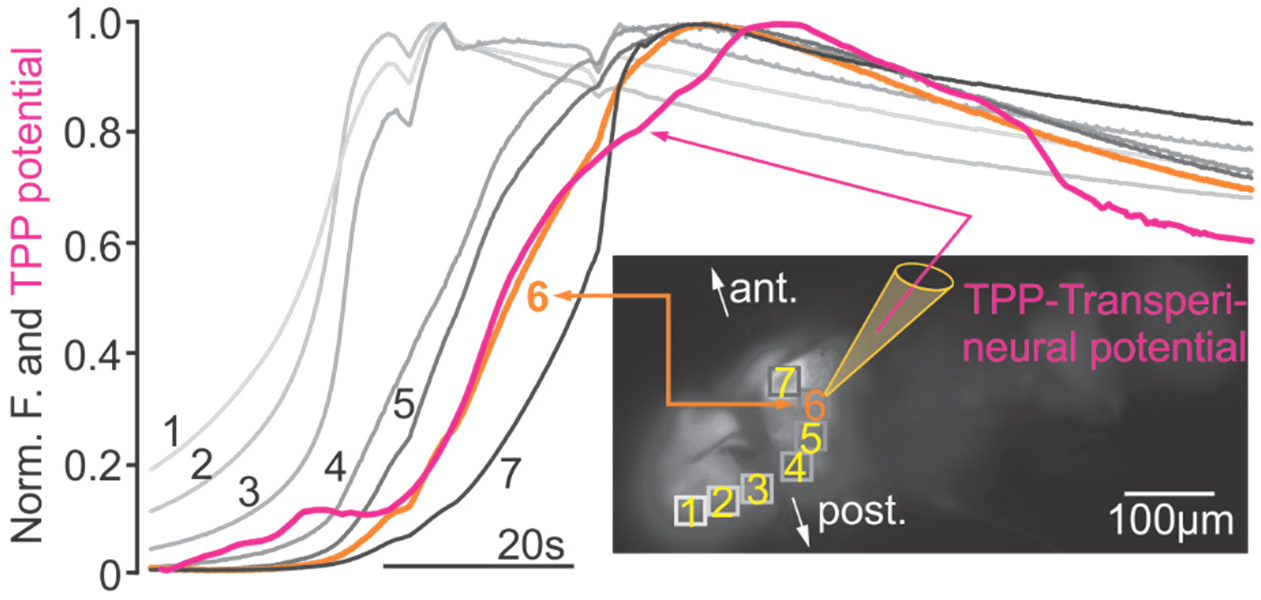
Original recording of GCaMP6m fluorescence during an SD event in an adult fly. Regions of interest (ROIs) 1-7 are shown along the path of SD spread. A sharp microelectrode was impaled at ROI 6 and the transperineural potential (TPP) was measured. Left: Normalized GCaMP fluorescence (gray lines) over time for ROIs 1-7. The trace for ROI 6 is highlighted in orange. The TPP measured from the sharp microelectrode located at ROI 6 coincided with the occurrence of the SD wave front as indicated by the change in fluorescence at ROI 6. For display purposes, the TPP was inverted. Ant. = anterior, post. = posterior.

Our results so far show that rapid cooling of the fly brain elicits a spreading wave of fluorescence consistent with the properties of SD and suggest that changes in the extracellular potassium concentration coincide with the wave. Because it is well-established that increasing the extracellular potassium concentration through application of potassium chloride (KCl) can elicit SD^31^, we performed calcium imaging of adult fly brains while we manipulated the extracellular potassium concentration at room temperature. We first increased the potassium concentration in the saline bath that surrounded the fly brain by spiking the bath with a drop (4µl) of 1M KCl solution. This rapidly increased the potassium concentration from the normal 5 mM to 12.9 mM. Our data show that in all cases (N=9) spreading waves of high intensity GCaMP6m fluorescence were observed. Fig. 5A shows an example recording where a wave was initiated in the right mushroom body and spread laterally. Shortly after, a second wave started in the left mushroom body, also spreading laterally. Finally, we observed a wave that traveled toward the calyx. This happened as the wave in the right mushroom body had already passed and fluorescence in the right hemisphere had started to diminish. The mean fluorescence over time plot of the whole brain (Fig. 5B) resembled those of the cooling experiments. The three wave occurrences are clearly visible in the fluorescence plot. When the wave began in the right mushroom body, a rapid rise of fluorescence occurred (peak at ii, orange). This was followed by the wave in the left mushroom body, which caused a subsequent steep rise in fluorescence (peak at iii, green). Lastly, the delayed wave near the calyx caused a smaller peak after the mean fluorescence had already reached its maximum (iv, red). Figure 5C shows the timing of the wave spreading across the brain, starting with the right mushroom body.

**Figure 5.**
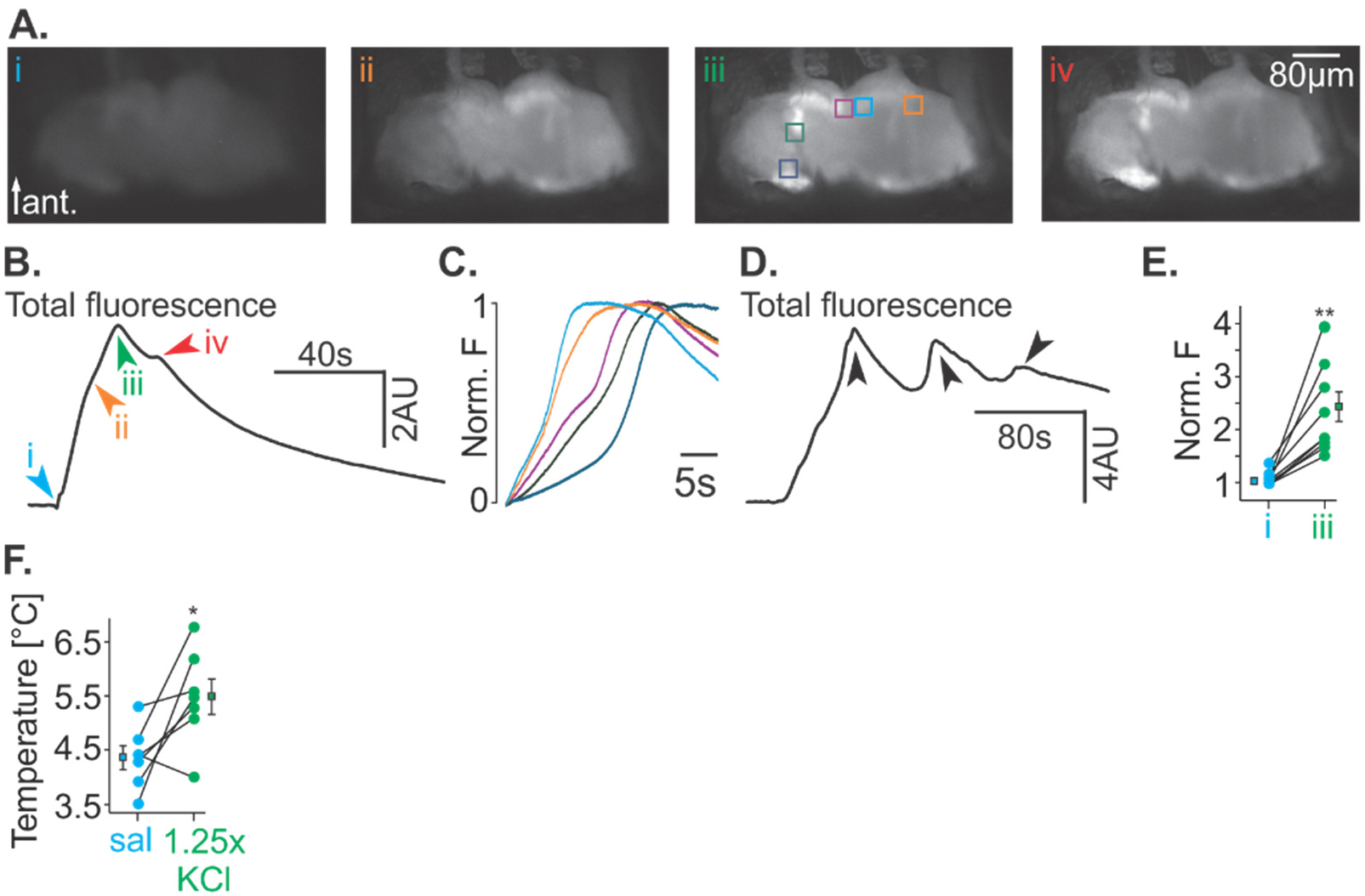
Increased potassium concentration facilitates the initiation of a spreading wave of calcium fluorescence. A) Representative original recordings of GCaMP6m fluorescence at room temperature while the brain was bathed in saline and then spiked with increased potassium. i, before KCl application; ii, shortly after KCl had been applied; iii, at peak fluorescence; iv, after fluorescence started to diminish. B) Whole brain mean fluorescence over time for the images shown in (A). C) Normalized mean fluorescence at the ROIs shown in panel Aiii. The minimum to maximum fluorescence was normalized to 0 to 1. D) Whole brain mean fluorescence changes over time in an animal that exhibited multiple waves of spreading fluorescence. E) Normalized mean fluorescence at the start of the spreading wave (i) and the end of the spreading wave when maximum fluorescence occurred (iii). The change in fluorescence was highly significant (N=9, P< 0.01, paired t-test). For each animal, mean fluorescence was normalized to the mean fluorescence value at the start of the experiment. F) Quantification of the temperature at which the spreading fluorescence wave was initiated in experiments where brains were cooled in standard physiological saline (sal) and in saline with enhanced (1.25x) KCl concentration. The initiation temperature in the KCl-enhanced saline was significantly higher than the temperature in standard saline (N=7, P<0.05, paired t-test).

In some experiments, multiple waves across the same brain areas were obtained, even though potassium was only applied once. Figure 5D shows an example where three independent waves were observed in response to a single KCl application. On average, the first spreading wave was observed within 14.0 ± 4.4 seconds of the application and reached a maximum fluorescence that was significantly larger than immediately before the wave started (before application: 1.0 ± 0.04, after application: 2.3 ± 0.3, paired t-test, P = 0.0015, N = 9; Fig. 5E).

In a second experiment, we tested the hypothesis that enriched extracellular potassium will facilitate the occurrence of SD, even at concentrations that are insufficient to elicit SD itself. For this, we bathed the brain in saline solution containing a 25% higher potassium concentration (6.25 mM) than standard saline (5 mM). For these experiments we first cooled the brain in regular saline to elicit a spreading wave of calcium fluorescence. After recovery to room temperature, we exchanged the saline with the enriched potassium saline and after 25 minutes, we repeated the cooling. We predicted that if potassium changes in the extracellular space are causally involved in eliciting the cooling-induced fluorescence wave, then bathing the brain in saline with elevated potassium levels should cause the rapid increase in fluorescence to occur earlier (i.e., at a higher temperature). Indeed, we found that on average, the temperature at which the rapid increase in fluorescence occurred was significantly higher in potassium enriched saline than in control saline (control saline 4.4 ± 0.2; potassium enriched saline: 5.5 ± 0.3, paired t-test, P = 0.03, N = 7; Fig. 5F).

Taken together, we have identified a slowly propagating wave of sustained neuronal depolarization that is induced by cooling or high potassium application, is modulated by extracellular potassium concentration, and is accompanied by changes in the extracellular potential. These observations are reminiscent of the hallmarks of SD, and thus consistent with the hypothesis that adult fly brains can exhibit SD.

### Cooling elicits a slowly spreading fluorescence wave in larval fly brains

While cooling-induced SD had previously been suggested to exist in adult fruit flies, it remains unknown whether cooling can also elicit SD in developing larvae, and if so, whether the developmental stage of the fly brain affects its susceptibility for SD. Existing literature suggests that SD results from underlying fundamental mechanisms that are shared between different circuits and brain structures (for review see^2^). We thus hypothesized that SD can also be induced in larval fruit flies and is based on the same mechanisms as those in the adult fly brain. To test this hypothesis, we mounted larval flies (first, second, and third instars) with pan-neuronal GCaMP6m expression on a cooling platform (see Fig. 6A and Material and Methods) and recorded calcium fluorescence of their entire nervous system. We employed two distinct mounting assays: In the first, our recordings were aided by the fact that the *Drosophila* larval stages are translucent, which allowed us to determine whether SD occurred in fully intact animals without the need to dissect out the nervous system (Figs. 6B, C). In the second assay, we used the ‘fillet’ dissection^32^ (Material and Methods) to open the dorsal cuticle and expose the nervous system (Figs. 6D,E). This allowed us to track fluorescence changes in more detail due to the exposed nervous system and to manipulate the saline solution surrounding the nervous system. The two assays together allowed us to monitor several animals simultaneously in a high-throughput approach and to do high resolution imaging of individual nervous systems.

**Figure 6.**
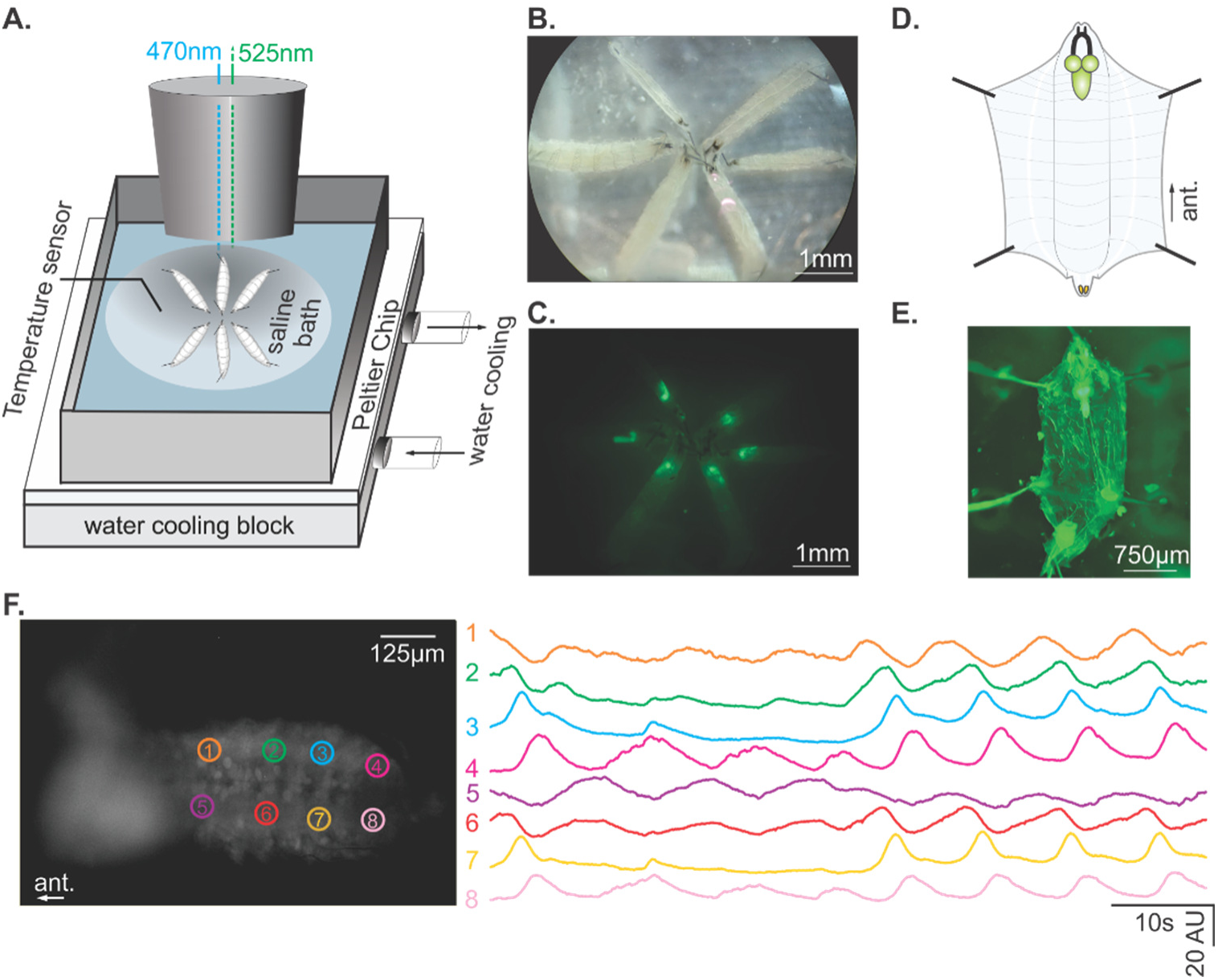
Whole nervous system fluorescent imaging of larval flies. A) Schematic of high-throughput experimental setup to rapidly cool larvae from room temperature to near 0℃ while measuring GCaMP6m or Arclight fluorescence in the nervous system. Up to 16 second instar larvae can be imaged simultaneously. B) Brightfield image of 6 second instar larvae. The anterior ends of the larvae with visible mouthhooks are pinned at the center and the posterior ends are pinned at the outer edge of the image. C) Same larvae as in (B), expressing pan-neuronal GCaMP6m under fluorescent light (470 nm illumination, 525 nm detection). The brain lobes and ventral nerve cords (VNCs) are clearly visible through the cuticle. D) Schematic of larval fillet dissection used for high-resolution imaging or for pharmaceutical applications. The anterior end of the larva was pinned at the top of the schematic and the posterior end was pinned at the bottom of the schematic. E) Fluorescence image of fillet-dissected larvae, dorsal view. The fluorescent brain and VNC are clearly visible after removal of the dorsal cuticle and gut. F) Spontaneous activity in the larval nervous system. Left – regions of interest (ROIs) used to measure spontaneous activity in theVNC. Right – fluorescence (arbitrary units) measured in each ROI at room temperature. Traces are separated vertically for clarity. See also supplemental 4.

Figure 6F shows a GCaMP6m fluorescence image of a single third instar nervous system at room temperature. Specifically, we focused on the ventral nerve cord to identify whether our approach allowed the detection of neuronal activity. As expected, and reported previously^33,34^, clearly visible slow waves of calcium fluorescence traveled posteriorly along the nerve cord. These waves were coordinated and had high signal-to-noise ratio (see also supplement 4), suggesting that they resulted from neuronal activity that drives larval crawling behaviors.

Like in adults, we cooled nervous systems at rates between 0.8°C/min and 3°C/min from room temperature (20 - 22°C) to near 0°C and then warmed back to room temperature at a rate of 3°C/min. Again, cooling ramps were kept consistent between trials for the same experiment, but different experiments tested different ramps (e.g., Fig. 7E). Figure 7A shows an example of the observed fluorescence in a third instar larva. Panel i shows the fluorescence after the brain had been cooled to 13°C. Panel ii shows the same brain after further cooling to 8.5°C, where a slight overall increase in fluorescence was observed (see also Fig. 7B). Panel iii shows a large increase in fluorescence at 6.1°C that was visible in the brain hemispheres and the ventral nerve cord. The brightest fluorescence occurred just before the coldest temperature of 0.9°C. Fluorescence decreased slowly during the initial re-warming phase (Panel iv; 8.1°C). Finally, the fluorescence decreased more rapidly when the brain was warmed back to 18°C (panel v). Figure 7B shows the mean fluorescence of the whole nervous system over the course of this experiment, starting at 18°C, cooling to 0.9°C, and then warming back to 18°C with the arrows and numbers indicating the panels shown in Figure 7A. Just as with adult brains, the changes in GCaMP6m fluorescence could be characterized into 4 regimes: 1) a slow global increase in fluorescence during initial cooling, 2) a rapid rise in fluorescence starting on average at 6.0 ± 0.3°C (N=30) across all larval stages, 3) a slow global decrease in fluorescence, and 4) a rapid decrease in fluorescence. This suggested that SD was present even in the larval fly nervous system. The observed changes in fluorescence occurred in all tested animals, regardless of larval stage. To characterize the effects of cooling on the different larval stages, we first compared results from experiments in which the same cooling rate (3°C/minute) was used. This avoided a potential influence of stimulus dynamics on the observed results. Figure 7C shows that the fluorescence change in all three larval stages was significant (N=10 animals of each instar, P<0.001 each, paired t-tests between start of fluorescence rise and maximum fluorescence). However, there were also differences: on closer inspection, we found that there was an overall significant effect of larval stage on SD initiation temperature (Fig. 7D, P<0.001, repeated measures Anova, N=10, F(2,18)=16.321) with the third instar larvae requiring a significantly colder stimulus to elicit SD than the first or second instar larvae (Student-Newman-Keuls posthoc test with P<0.05). Specifically, in these experiments, we paired all three larval stages on the same cooling plate to directly compare the effects of cooling between developmental stages.

**Figure 7.**
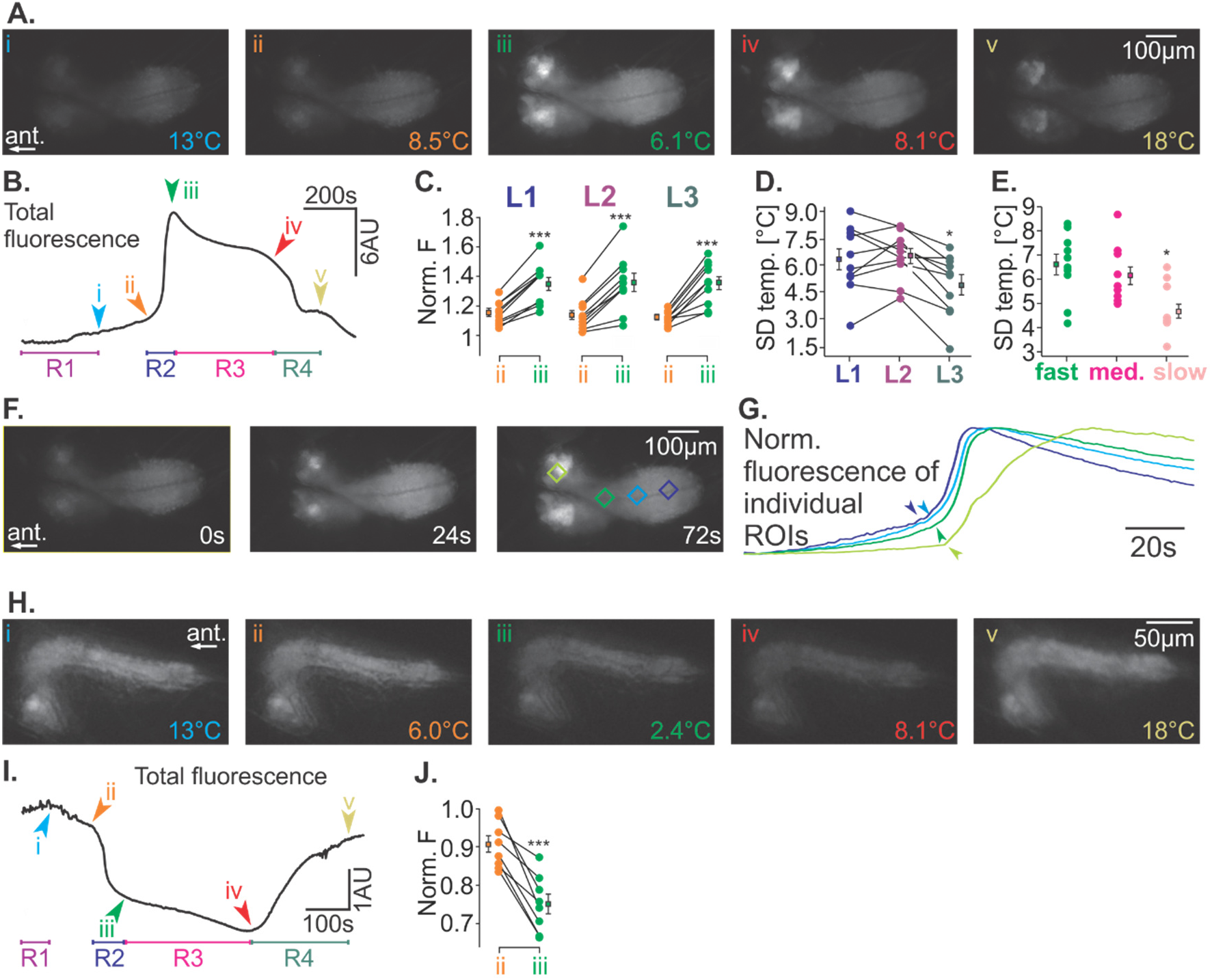
Changes in GCaMP6m fluorescence during cooling and rewarming of the larval nervous system can be classified into the same 4 regimes the adult brain. A) Representative original recording of GCaMP6m fluorescence during cooling to near 0℃ and then rewarming back to room temperature. i) End of regime 1, 13℃. ii) Start of regime 2 at 8.5℃. iii) End of regime 2 = start of regime 3 at 6.1℃. iv) End of regime 3 = start of regime 4 at 8.1℃. v) End of regime 4, at 18℃. B) Representative mean fluorescence of the whole nervous system over time showing the 4 distinct regimes (R1 – R4) and their starting and ending points (i – v). C) Quantification of normalized mean fluorescence at points ii (start of the quick fluorescence rise) and iii (peak fluorescence). There was a significant increase in fluorescence in all larval stages (instars 1, 2, and 3; separate paired t-tests, P<0.001 each). For each animal, mean fluorescence was normalized to the mean fluorescence value at 18 °C at the start of the cooling ramp. D) Comparison of SD initiation temperature in the three larval stages. The third instar larvae required significantly colder temperatures to elicit SD (repeated measures Anova, N=10, F(2,18)=16.321, Student-Newman-Keuls posthoc test with P<0.05). E) Comparison of different cooling speeds on SD initiation temperature (slow: 0.8°C/minute, medium: 1.5°C/minute, fast: 3°C/minute). The SD temperature of the slowest cooling rate was significantly lower (overall effect of cooling P=0.002, one way Anova, N≥10, F(2,28)=7.764), with Student-Newman-Keuls posthoc test with P<0.05). F) Original recordings of GCaMP6m fluorescence during regime 2 showing that the fluorescence wave spreads from the VNC to the brain hemispheres. Times of the frames are shown from the start of the spreading wave (t = 0). The right image shows 4 regions of interest (ROIs) used to measure the fluorescence in (G). See also supplemental 5. G) Normalized fluorescence for the ROIs shown in (F). The arrows indicate the start of regime 2 and the start of the SD event for each ROI. They are clearly separated in time, indicating that the fluorescence wave shows a spatial spread. H) Same as (A), but for Arclight fluorescence. Lateral view of the nervous system. Temperatures at each point are indicated in the figure. I) Same as (B), but for Arclight fluorescence in second instar larvae. J) Quantification of normalized mean fluorescence at points ii (start of the quick fluorescence drop) and iii (trough of fluorescence). There was a significant decrease in fluorescence (paired t-test, P<0.001).

Finally, we also tested the effects of different cooling rates on SD temperature. Second instar larvae were exposed to either 0.8°C/minute, 1.5°C/minute, or 3°C/minute cooling ramps. We found a significant effect of cooling rate on SD temperature (Fig. 7E; P=0.002, one way Anova, N≥10, F(2,28)=7.764), with the slowest cooling rate requiring significantly colder temperatures to elicit SD (Student-Newman-Keuls posthoc test with P<0.05).

To confirm that the observed fluorescence changes were caused by a spreading wave, we quantified the fluorescence change at several points throughout the nervous system. Figures 7F,G show the result of this analysis, where fluorescence started to increase in the posterior section of the ventral nerve cord, and then slowly spread toward the anterior brain hemispheres (see also supplemental 5). We found that between animals the waves differed in starting location but were typically either initiated in the anterior brain lobes or in the posterior third of the ventral nerve cord. From their initiation point, waves spread throughout the entirety of the larval nervous system. Interestingly, we found that on average, spread velocities were slower than those in the adult brain (6.1 ± 0.6µm/s, N=12, P=0.049, t-test).

To confirm that the observed waves in GCaMP6m fluorescence were due to increased neuronal membrane depolarization, we again used a fly line with pan-neuronal expression of the voltage sensor Arclight and repeated the cooling experiment. Similar to adult brains, we found that when cooled from room temperature, there were initially small and slow changes in fluorescence, and that the overall Arclight fluorescence response was inverted from that of GCaMP6m. The initial slow changes in Arclight fluorescence were followed by a rapid drop in fluorescence, suggesting that larval neurons underwent a rapid depolarization. Figure 7H shows an original recording of such an experiment, with panel i showing the mean fluorescence at 13°C. There was a slight drop in fluorescence as cooling continued (panel ii; 6.0°C), followed by a larger and quicker drop in fluorescence in panel iii (2.4°C). Upon warming, the fluorescence continued to slowly decrease (panel iv; 8.1°C), but then rapidly increased as the temperature returned to 18°C (panel v). Fig. 7I shows the total fluorescence obtained during this trial across the four regimes. On average, the largest drop in fluorescence occurred at an average temperature of 5.3 ± 0.9°C (N=8) and reached its minimum at 3.0 ± 0.7°C (N=8). The lowest fluorescence was significantly different from the initial fluorescence (paired t-test, P<0.001; Fig. 7J). Taken together, our Arclight and GCaMP6m data thus suggest that larval fly neurons resemble those in adult brains during cooling and that they undergo the physiological changes that have been described for SD^2^).

To test whether the spreading fluorescent wave observed in larvae can also be initiated by increasing the extracellular potassium concentration, we used the fillet dissection with pan-neuronal GCaMP6m expression. Like in adults, we rapidly increased the potassium concentration in the saline bath surrounding the brain by spiking the bath with a 1M KCl solution. Several animals were mounted in each experiment (range: 2 - 4), and in all animals (N=12), we observed a spreading wave of rapidly increasing fluorescence that was initiated within 3.7 ± 1.3 seconds of the application. Figure 8A shows an example recording where a wave was initiated in the left brain lobe and spread posteriorly to the ventral nerve cord, but also laterally to the right brain lobe. Panel i shows the fluorescence before KCl was applied. The image in panel ii was taken shortly after the fluorescence wave started. Panel iii shows the peak fluorescence when the wave had reached the ventral nerve cord. Finally, panel iv shows the slightly diminished fluorescence after the wave had spread throughout the whole nervous system, and brain lobe fluorescence started to diminish. The mean fluorescence over time plot of the whole nervous system is shown in Figure 8B. It closely resembled that of the cooling experiments, with a rapid rise in fluorescence shortly after KCl application. Figure 8C shows the timing of the wave spreading across the brain, starting with the left brain lobe (blue). The wave then reached the medial section of the ventral nerve cord (orange), before spreading into the right brain lobe (purple) and the posterior ventral nerve cord (green). We observed spreading waves of fluorescence in all tested animals (N=12). On average, the fluorescence peak was 5.1 ± 0.3 times higher than before KCl application (Fig. 8D; P<0.001, paired t-test, N=12).

**Figure 8.**
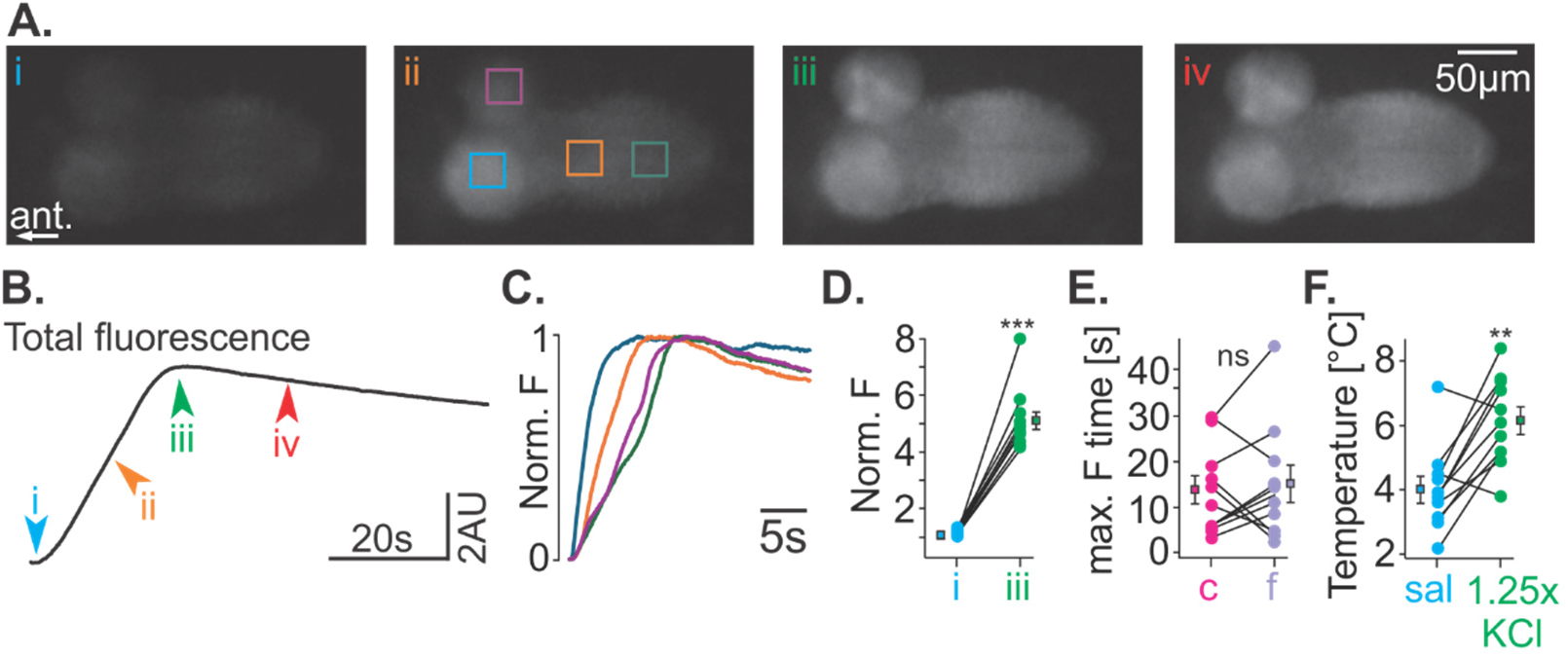
Increased potassium concentration facilitates the initiation of SD in the larval nervous system. A) Representative original recordings of GCaMP6m fluorescence at room temperature while the nervous system was bathed in saline and then spiked with increased potassium. i, before KCl application; ii, shortly after KCl had been applied; iii, at peak fluorescence; iv, after fluorescence started to diminish. B) Whole nervous system mean fluorescence over time for the images shown in (A). C) Normalized mean fluorescence at the ROIs shown in panel Aii. The minimum to maximum fluorescence was normalized to 0 to 1. D) Normalized mean fluorescence at the start of the spreading wave (i) and the end of the spreading wave when maximum fluorescence occurs (iii). The change in fluorescence was highly significant (N=12, P< 0.01, paired t-test). For each animal, mean fluorescence was normalized to the mean fluorescence value at the start of the experiment. E) Peak mean fluorescence time at the closest and farthest points from the location of the KCl application were not significantly different (N=10, paired t=test, P=0.70). F) Quantification of the temperature at which the spreading fluorescence wave was initiated in experiments where nervous systems were cooled in standard physiological saline (sal) and in saline with enhanced (1.25x) KCl concentration. The initiation temperature in the KCl-enhanced saline was significantly higher than the temperature in standard saline (N=11, P=0012, paired t-test).

Since we noticed that in most cooling experiments, the fluorescence waves either started near the anterior brain lobes or at the posterior end of the ventral nerve cord, we were curious whether the location of the KCl application would bias the initiation site. In our high-throughput approach, we monitored several larvae simultaneously while they were pinned in different orientations (e.g., Fig. 6A). We then determined the time of peak fluorescence at the closest and farthest points of the nervous system from the location of the KCl application. Peak times were measured from the time of the KCl application. Figure 8E shows that there was no significant difference between peak times, indicating that the location of the drop relative to the nervous system did not bias spread initiation (N=10, paired t=test, P=0.70).

Finally, we tested the hypothesis that enriched extracellular potassium also facilitates the occurrence of SD in larval fly brains. For this, we bathed second instar nervous systems in saline solution containing a 25% higher potassium concentration (6.25 mM) than standard saline (5 mM). This concentration was insufficient to elicit SD at room temperature. As a control, we first cooled the nervous system in regular saline and measured the temperature at which SD occurred. After recovery to room temperature, we exchanged the saline with the enriched potassium saline and after 25 minutes, we repeated the cooling. Based on our results in adult brains (Fig. 5F), we predicted that SD would occur earlier (i.e., at a higher temperature). Indeed, we found that on average, SD temperature was significantly higher in potassium enriched saline than in control saline (control saline 4.0 ± 0.4; potassium enriched saline: 6.2 ± 0.4, paired t-test, P = 0.0012, N = 11; Fig. 8F). This suggests that potassium changes in the extracellular space are causally involved in eliciting the cooling-induced SD. Taken together, our data thus indicated that the larval fruit fly nervous system reliably experienced spreading depression both during cooling and after high potassium application.

### Spreading depression confers protection against subsequent spreading depression

Are there potential lasting effects of spreading depression on future events? To address this question, we designed an experiment with repeated cooling cycles. Specifically, we imaged second instar brains with pan-neuronal expression of GCaMP6m and a 3°C/min cooling ramp. Five cooling and rewarming cycles were used, with a 5-minute break in between each cycle (approximately 15 minutes total per cycle). In each cycle, we measured the temperature at which SD occurred, and then compared these temperatures across the five cooling cycles. We found that there was an overall significant effect of the cooling cycles (P<0.001, repeated measures Anova, N=8, F(4,28)=21.341). The SD temperatures of cycles 1, 2, and 5 were significantly different from all other cycles (Fig. 9A), with cycle 1 being the easiest to elicit (having the highest SD temperature: 7.5 ± 0.3°C, N=8) and cycle 5 being the hardest to elicit (having the lowest SD temperature: 3.6 ± 0.7°C, N=8). The SD temperatures of cycles 3 and 4 were not different from one another, but they were significantly different from all other cycles (Student-Newman-Keuls posthoc test with P<0.05).

**Figure 9.**
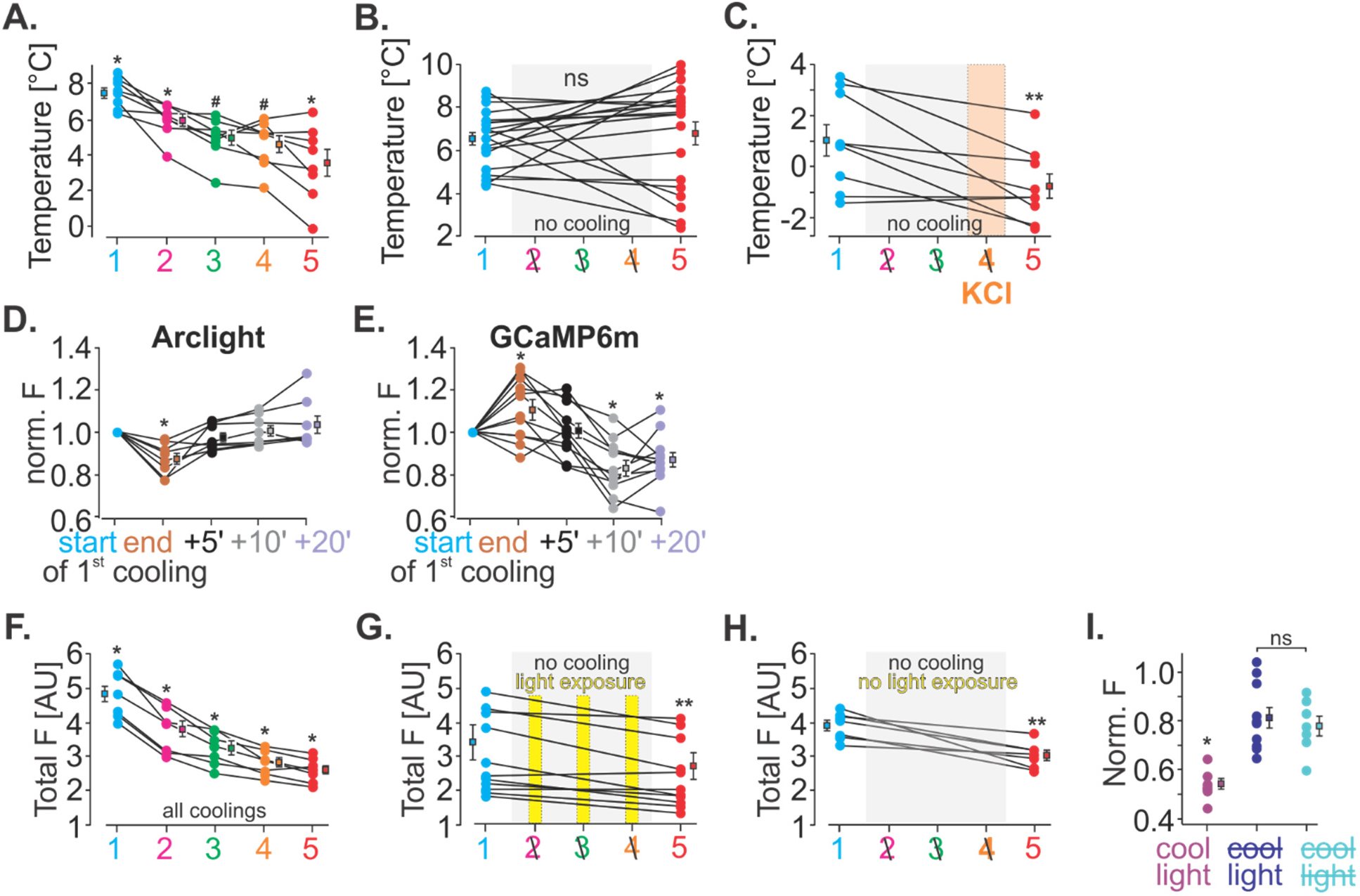
SD confers short-term resilience to subsequent SDs (for detailed statistics, see text). A) Quantification of SD temperature of multiple SDs that were elicited by repeated cooling of second instar nervous systems. Subsequent SDs required colder temperatures to elicit SD. B) With sufficient time between cooling cycles, there was no significant effect on SD temperature. Only cooling cycles 1 and 5 were applied. C) When SD was elicited with KCl application to the nervous system, the effect was re-established, and the subsequent cooling-induced SD required a colder temperature to be elicited. Only cooling cycles 1 and 5 were applied. KCl was applied instead of cooling cycle 4. D) Quantification of normalized Arclight fluorescence at room temperature before the first cooling cycle, and the end of the first cooling cycle, and 5, 10, and 20 minutes later. Only the fluorescence at the end of the cooling cycle was significantly lower. E) Quantification of normalized GCaMP6m fluorescence at room temperature before the first cooling cycle, and the end of the first cooling cycle, and 5, 10, and 20 minutes later. F) Whole nervous system mean fluorescence across 5 cooling-induced SD events. Fluorescence was measured at the start of each cooling cycle. There was a significant drop in fluorescence with each SD. G) Whole nervous system fluorescence when only the first cooling-induced SD was elicited. While cooling cycles 2 – 4 were not applied, and no SD was elicited, the fluorescence light was turned on for the corresponding time. Fluorescence was significantly lower before cooling cycle 5. H) Whole nervous system mean fluorescence when only the first cooling-induced SD was elicited. Here, the fluorescence light was kept off after the first cooling. Fluorescence was still significantly lower before cooling 5. I) Comparison of whole nervous system fluorescence before cooling cycle 5 from experiments shown in E-F, normalized to the fluorescence before the respective first cooling trials. Fluorescence was significantly lower when repeated SDs were elicited. Having the fluorescence light on did not significantly affect the measured fluorescence.

While these data suggested that a single SD event can influence the likelihood of subsequent ones, we wanted to exclude a potential effect of the duration of the experiment on our results. We thus altered the experimental approach such that instead of 5 repeated cooling cycles, we only carried out the first one, then skipped cycles 2 through 4 by leaving the animal at room temperature, and finally carried out cooling cycle 5. We kept the time between cooling cycle 1 and 5 the same as in the original experiment (3 x 15 minutes = 45 minutes), i.e., cycle 5 was started at its usual time 60 minutes after the start of cycle 1. In this case, when we compared SD temperatures between cooling cycles 1 and 5, we no longer found a significant difference (Fig. 9B; N=20, paired t-test, P=0.65).

Finally, to confirm that the effect of repeated cooling cycles on SD temperatures was due to the SD event itself and not the exposure to cold temperature, we repeated the experiments where we skipped cooling cycles 2 to 4. However, at the time where cooling cycle 4 was expected (45 minutes after cycle 1 started), we elicited SD through application of 1M KCl to the bath (similar to our previous experiments, see Fig. 8). SD was observed in all animals tested. After the KCl-elicited SD occurred, KCl was washed out with regular saline, and the fifth cooling cycle was started at its usual time (60 minutes after the start of cycle 1). We found that the SD temperature of cycle 5 was again significantly lower than that of cycle 1 and thus SD was harder to elicit (Fig. 9C; N=9, paired t-test, P=0.0093). Overall, this suggests that the initial cooling-induced SD event provided long-lasting effects that counteracted the initiation of subsequent spreading depression events, and that a single event was sufficient to cause that effect.

Because these experiments were carried out in second instar larvae, we wondered whether the observed resilience mechanisms occur at other developmental stages. We thus carried out repeated cooling cycles in first and third instar larvae, as well as in adult flies (supplemental 6). In all cases, we found an overall significant effect of the repeated cooling cycles, with the SD temperature of the first cycle being the significantly higher than all other cycles in all tested groups. In all groups, the SD temperatures of cycles 2 and 5 were significantly lower than that of cycle 1. Specifically, in first instar larvae, there was an overall significant effect of cooling cycle (P<0.001, repeated measures Anova, N=7, F(6,24)=24.169). The SD temperature of cycle 1 was significantly higher than those of all other cycles. The SD temperature of cycle 2 was significantly higher than all subsequent cycles, and the SD temperature of cycle 3 was significantly higher than that of cycle 5, but not of cycle 4 (Student-Newman-Keuls posthoc test with P<0.05). In third instar larvae there was an overall significant effect of the cooling cycle (P<0.001, repeated measures Anova, N=9, F(4,32)=6.843) and the SD temperature of cycle 1 was significantly higher than in all other cycles (Student-Newman-Keuls posthoc test with P<0.05). Finally, in adults, there was an overall significant effect of the cooling cycle (P<0.001, repeated measures Anova, N=8, F(4,28)=6.548). The SD temperature of cycle 1 was significantly higher than those of cycles 2 and 5 and the SD temperatures of cycles 3 and 4 were significantly higher than that of cycle 5 (Student-Newman-Keuls posthoc test with P<0.05).

To test whether we could identify a potential lasting effect of repeated SDs on the activity of the neurons, we again turned to second instar larvae, and imaged fluorescence with pan-neuronal Arclight and GCaMP6m expression. In both cases, we saw that in most recordings, the fluorescence levels after cooling did not fully return to baseline levels from before cooling (e.g., Fig. 7). We thus compared the Arclight and GCaMP6m fluorescence over extended periods of time at room temperature after the first cooling-induced SD. For Arclight, we found that immediately after the cooling (i.e., at the end of the cooling ramp), the fluorescence was significantly lower than at the beginning of the cooling trial (Fig. 9D; repeated measures Anova, N=8, F(4,28)=12.506, Student-Newman-Keuls posthoc test with P<0.05 calculated with unnormalized data). However, 5 minutes later, Arclight fluorescence had returned to baseline levels and was no longer significantly different from before the trial. This was also the case after 10 and 20 minutes, suggesting that SD had no lasting effect on the activity of the neurons. For GCaMP6m fluorescence, however, we found that immediately after the cooling the fluorescence was significantly higher than at the beginning (Fig. 9E; repeated measures Anova, N=12, F(4,44)= 12.717, Student-Newman-Keuls posthoc test with P<0.05 calculated with unnormalized data), returned to baseline 5 minutes after the cooling cycle, but then was significantly lower after 10 and 20 minutes. Also, subsequent repeated cooling cycles (and thus SDs, Fig. 9F) resulted in significantly reduced GCaMP6m fluorescence levels. Specifically, there was an overall significant effect of the repeated cooling cycles, and the fluorescence at the start of each successive cooling cycle decreased significantly (P<0.001, repeated measures Anova, N=8, F(4,28)=121.18, Student-Newman-Keuls posthoc test with P<0.05).

To test whether this lasting change in calcium fluorescence was due to the repeated spreading depression events or was a consequence of the long experimental time and continuous exposure to the blue excitation light used for imaging, we carried out two control experiments. First, instead of 5 repeated cooling cycles, we only carried out the first one, then skipped cycles 2 through 4 before cooling as usual during cycle 5. During skipped cooling cycles, the animals were left at room temperature for a total of 45 minutes (i.e., the time corresponding to cooling cycles 2-4) and were exposed to the excitation light as usual for trials 2 - 4. Despite the absence of cooling, we found a significant decrease in GCaMP6m fluorescence after 45 minutes (Fig. 9G, paired t-test, N=11, P=0.0015), indicating a potential effect of the experiment duration or the continued light stimulus on either the GCaMP6m proteins or the calcium levels in the neurons. We thus carried out a second control experiment in which we repeated the first control experiment, but did not expose the animals to excitation light or cooling for cycles 2 - 4. This still resulted in a significant reduction in fluorescence over time (Fig. 9H, paired t-test, N=7, P=0.003). Taken together, this suggested that the mere long duration of the experiment led to a diminishment of the GCaMP6m fluorescence. We thus wondered whether this effect could explain the reduction in fluorescence we had observed with each successive SD events (Fig. 9F). To test this, we measured the relative change in fluorescence between the start of cooling cycle 1 and the start of cooling cycle 5. We then compared experiments where we repeated all cooling cycles with those where only the first cooling cycle was carried out and animals were exposed to repeated light stimuli but no further cooling cycles, and those where only the first cooling cycle was carried out and animals were not exposed to repeated light stimuli or further cooling cycles (Fig. 9I). We found that the experiments where repeated cooling cycles were applied showed a significantly larger reduction in fluorescence than the two experiments in which no cooling was applied (One-way Anova, 7≤N≤11, F(2,23)= 16.259, Student-Newman-Keuls posthoc test with P<0.05). In addition, the two experiments without cooling were not different from one another. Collectively, these experiments thus demonstrate that while there was a small effect of time on our fluorescence signal, repeated SDs significantly altered the baseline calcium fluorescence in larval neurons.

Taken together, our data show that cooling-induced SD events provide a long-lasting effect that counteracts the initiation of subsequent SD events, and that these effects act independently of the brain’s developmental stage. Our experiments also demonstrate that the mechanisms providing this resilience are temperature-independent and directly activated by the occurrence of SD.

## III. Discussion

We demonstrate that spreading depression can be observed in whole brain calcium and voltage imaging in adult and larval *Drosophila melanogaster*. SD could either be elicited through rapid cooling, a naturally occurring environmental condition, or through application of high potassium chloride. We identified four regimes of our calcium and voltage imaging data, including an initial slow rise followed by sudden increase in membrane voltage, sustained depolarization, and rapid recovery. These regimes correspond to characteristic features of SD observed in other systems, including the rapid and dramatic changes in neuronal physiology observed in mammals and insects. In particular, SD is associated with an increased firing frequency that leads to depolarization block^2,8,35^ along with significant alteration of ion concentrations and gradients across the neuronal membrane.

Prior studies had provided evidence of a putative SD in insects, including locusts, butterflies, and *Drosophila*. These studies measured a sharp decrease in extracellular potential at two spatially and temporally separated points when the adult *Drosophila* nervous system was rapidly cooled, but it remained unclear whether the observed changes at the two points were connected rather than separate events. Our results definitively confirm through calcium and voltage imaging that a slowly travelling wave of depolarization spreads across the adult *Drosophila* brain and that the occurrence of this wave coincides with the decrease in extracellular potential. Through SD threshold measurements in elevated potassium saline, we have shown that the process is potassium-dependent and that increased extracellular potassium facilitates SD initiation.

In adult fly brains, we observed SD initiation at multiple sites with a singular stimulus, and in our experiments with repeated cooling cycles, we observed that SD could initiate in different locations with each repeated stimulus. This indicates that SD does not result from a defect in a singular location and that it can be initiated at many sites. SD initiation is thus not specific to a given neuronal architecture, connectivity, or cell type population. However, we observed a high prevalence for SD spreading through the mushroom body, perhaps indicating that this region is more prone to SD or that it is more readily observable here. The mushroom body is composed of about 2000 Kenyon cells^36,37^ with densely packed cell bodies near the calyx and axons projecting anteriorly into the pedunculus before splitting into the alpha, beta, and gamma lobes. Our observations showed that SD spread typically began near the calyx of one hemisphere before spreading to the pedunculus and alpha lobe, followed by the beta and gamma lobes of that hemisphere. In many cases, it then jumped to the gamma and beta lobes of the other hemisphere and spread in a reverse pattern from the alpha lobe and pedunculus and finally into the calyx. This pattern suggests that SD propagates through sites with high cell body density and sites with many axons, but also that it is mediated through local changes to the region experiencing SD. If instead SD spread was mediated through electrical signaling in the axons (i.e., action potential propagation), one would expect a rapid jump from the initiation site at the calyx to the site where axons terminate (the mushroom body lobes). However, this was not observed and the fluorescence wave moved slowly through the MB into adjoining regions, consistent with the idea that SD spreads through diffusion of extracellular potassium. We also observed spread in other neuropils, including the superior lateral protocerebrum, the lobula, and the antennal lobes. The neurons in these structures serve vastly different functions, including visual^38^ and olfactory processing^39,40^, they are differently organized than those of the MB, and neuronal densities and synaptic connectivity differ. The occurrence of SD in all these areas thus indicates the universality of SD,

In mammals, there is evidence that SD does not cross white matter, preventing spread into different brain regions^2^. Likewise, in insects, there is evidence that SD does not cross between ganglia. Specifically, SD was shown to not spread between the thoracic ganglia of adult locusts^41^. However, our data shows that SD spreads between the brain lobes and VNC (or vice versa) of larval *Drosophila*. The larval nervous system consists of the brain and the VNC, which is a single consolidated ganglion, containing distinct and segmentally organized neuromeres (three fused subesophageal neuromeres, three thoracic neuromeres, seven abdominal neuromeres, and three terminal neuromeres). There is no clear spatial separation between brain and the VNC like in many adult insects, suggesting again that local diffusion, rather than the propagation of neuronal activity in axons underlies SD spread.

In addition to demonstrating that SD exists in adult flies, we show for the first time that SD exists in all three larval instars with properties similar to those of the adult brain. Structure, neuronal connectivity, and neuronal numbers are quite distinct in each of the three larval stages and the adults. The presence of SD across these developmental stages and its similarity between them thus points to SD being a fundamental phenomenon that occurs independently of these features. This idea is supported by the fact that SD occurs in the brains of animals from many different clades (including mammals, birds, and insects).

Despite the presence of SD across developmental stages, our results indicate several differences between larval and adult SD, each of which may help identify characteristics relevant to SD. In particular, the fold-change of fluorescence and the velocity of the spread was different in larvae compared to adults. In larvae, the fold change in calcium fluorescence was less than that of adults and the velocity of the spreading wave was slower in larvae compared to adults. These findings suggest that the overall excitation is lower in larvae than it is in adults. One contributing factor may be that the ratio of spiking to nonspiking neurons changes during development. *Drosophila* have only one type of sodium channel (*para*^42^), but with many isoforms. In third instar larvae, para is only expressed in 23% of the neurons^42^, suggesting that only this subset is able to fire Na_v_-dependent action potentials. In contrast, the adult central nervous system widely expresses para and therefore many more neurons may be able to fire action potentials^42^. If the accepted working hypothesis of SD is correct, and neurons undergo a rapid increase in firing frequency followed by depolarization block^8,35^, then it should be expected that a nervous system with fewer spiking neurons would behave differently than one with many. Specifically, in the system with fewer spiking neurons (e.g. larvae), there are fewer neurons to be trapped in depolarization block resulting in a lower overall average depolarization and delayed or reduced disruption of ion homeostasis. Because calcium fluorescence depends on the average depolarization of spiking and nonspiking neurons, these effects should manifest in a reduced fluorescence. Fewer spiking neurons in larvae may also explain the reduced spread velocity. SD is hypothesized to spread due to increased extracellular potassium and its diffusion to neighboring sites. With fewer neurons in depolarization block, potassium accumulation in the extracellular space would be predicted to be slower, with delayed buildup at neighboring sites. Despite the differences in numbers of spiking neurons, we were consistently able to elicit SD in both larvae and adults and did not see a difference in the SD initiation threshold. Additionally, we confirmed that in both larvae and adults, elevated extracellular potassium levels are causally related to SD initiation with a 25% increase in extracellular potassium leading to lower SD thresholds in both cases. Thus, the reduced number of spiking neurons in larval flies does not affect the occurrence or initiation of SD, nor its dependence on extracellular potassium.

Our results also show that a single SD event creates a protective effect which raises the threshold of future SD events. This protective effect was independent of how the SD was elicited, since both cooling and KCl application were able to establish it. The protective effect was present within a few minutes of the SD but was absent 45 minutes later. These results point to a cellular- or circuit-level memory trace from a lingering adaptation of the nervous system that at least partly protects against SD. Lasting effects have been observed in SD-related pathologies previously. In migraine disorders in humans, there is a 24-48 hour postdrome phase following the migraine, during which patients experience lasting symptoms, including fatigue, brain fog, and sensory sensitivity. However, patients also report a reduced likelihood for subsequent migraines during the postdrome phase, suggesting that the initial migraine may confer a temporary protective effect. While it remains unknown whether this protective effect is related to SD, lasting effects have been suggested even in the initial reports of SD^43^, and short-term protective effects reminiscent of what we observed in flies have been reported in mammalian and avian models of SD. In these cases, a refractory period lasting several minutes was observed, during which there was an increased threshold to elicit SD^27,44^.

In order for a lasting memory trace to exist, the initial SD event must cause changes that occur quickly and remain for many minutes. One such possibility is that the intra- or extracellular ion concentrations do not return to their pre-SD levels following the event. The massive redistribution of ions that occurs during SD includes a very large rise in extracellular potassium, and accompanying large drops in extracellular sodium, chloride, and calcium^2^. In our KCl application experiments, we saw multiple successive SD events, suggesting that extracellular potassium concentrations had recovered to the point where depolarization block was absent, and a new SD event could be initiated. In our fluorescence recordings, this is indicated by the rapid drop of calcium fluorescence after SD. It seems reasonable to assume that if the extracellular potassium concentrations remained abnormally elevated for minutes after the SD event, and the potassium equilibrium potential remained depolarized, this would lead to *<u>increased</u>*cell excitability and a *<u>lower</u>* threshold for initiating the next SD. We observed the opposite, suggesting that continued elevated extracellular potassium levels are unlikely to contribute to the observed memory trace.

However, altered extracellular potassium concentrations have long been known to activate homeostatic processes that cause lasting changes to neuronal activities and responses. For example, sustained exposure of rat hippocampal pyramidal neurons to elevated extracellular potassium led to persistent changes in neuronal excitability that were mediated by a calcium-dependent process that altered membrane potential and input resistance^45^. Similarly, long-term exposure of rat myenteric neurons to high extracellular potassium caused long-lasting alterations in calcium channel function^46^. These persistent effects are reminiscent of activity-dependent homeostatic plasticity that acts through changes in gene expression to restore cell excitability over many hours and days (e.g., ^47,48^). They are thus unlikely to have contributed to the observed memory trace after SD. Correspondingly, in studies of mammalian cortex pyramidal cells, only small changes to the excitability of neurons were observed, such as a mildly increased rheobase. However, no changes to input resistance or frequency-current (F/I) curves were present, suggesting that there are only minor alterations to the intrinsic properties of the neurons following SD^27^. In contrast to these findings, there is evidence for rapid and short-term (seconds to minutes) neuronal excitability changes after exposure to elevated extracellular potassium in crustacean stomatogastric neurons^49^. While the specific mechanisms remain unclear, it has been speculated that cell excitability is altered by rapid phosphorylation of voltage-gated ion channels. Additionally, the SD-induced changes in extra- and intracellular potassium (and sodium) concentrations are likely to lead to strongly activate Na^+^/K^+^ pump activity, in particular when temperatures recover from a cold-induced SD. Increased pump activity is known to lower cell excitability by hyperpolarizing the membrane potential. Such effects have been seen during bursts of high neuronal firing where pump activity is a contributor to spike frequency adaptation and increased after-hyperpolarizations that outlast burst activity for several seconds^50–52^. Indeed, our Arclight experiments suggest that there is a sustained hyperpolarization after the end of SD. They also suggest that this sustained hyperpolarization disappears within 5 minutes. In contrast, our calcium imaging results imply a longer-lasting reduction of neuronal excitability (> 30 minutes). These results are consistent with modeling studies of *Drosophila* larval neurons that suggest that Na^+^/K^+^ pump mediated reductions of cell excitability can outlast the rather short effects of the pump on membrane potential and persist in the absence of membrane potential changes^52^. These effects are predicted to be mediated by the interdependent actions of ion concentration changes, diffusion, and pump activation.

Lastly, the neuronal hyperexcitation during SD can lead to excessive release of neurotransmitters and cause lasting changes in neurotransmitter release, including glutamate and GABA^2^. Sawant-Pokam et al.^27^ provide evidence of altered transmitter release and presynaptic neuron firing in the mouse somatosensory cortex. This results in a sustained inhibitory shift in the ratio of excitatory to inhibitory synaptic inputs in layer 2/3 pyramidal neurons that lasted for up to one hour. It is conceivable that similar effects are at work in flies as well, leading to the protective effect we observed after SD. Such lasting effects could be mediated by synaptic plasticity (e.g., depression) in response to the excessive synaptic release during SD, or by enhanced and prolonged enzyme degradation of the released transmitters and facilitated transport into glial cells such as astrocyte-like glia or sub-perineuronal glia. Astrocyte-like glia are in intricate proximity to the neuropil and responsible for neurotransmitter clearance, e.g., glutamate uptake through the Excitatory Amino Acid Transporter (EAAT^53^). Sub-perineuronal glia regulate transport from the hemolymph into the ganglion, acting as a blood-brain barrier^54^. They hypertrophy during the third instar, which may explain why in the comparison between the different larval stages, the third instar larvae were the most difficult in which to elicit SD.

In all, our results demonstrate that SD is a phenomenon that persists across developmental stages and in networks with vastly different sizes, connectivity, and neuronal populations. Its occurrence in *Drosophila* shows similar characteristics to that of mammals, with slow spread across large brain areas and dramatic changes in neuronal behavior. We have shown that SD is facilitated by increased extracellular potassium, can originate from multiple independent sites, and that these sites can vary with repeated events. A singular SD event causes lasting effects observable in the depressed baseline calcium signal of the neurons and offers a limited-term protection from future SD events.

## IV. Materials and Methods

### Fly Stocks and Maintenance

Fly stocks were obtained from the Bloomington Drosophila Stock Center (Bloomington, IN). Flies were reared and maintained at room temperature (22° C) with the natural daily light cycle in vials with food (Formula 4-24 Instant Drosophila Medium, Plain) from Carolina Biological Supply Company (Burlington, NC). The first filial 1 (F1) generation was obtained by crossing virgin females of an nSyb-GAL4 line, w[1118]; P{y[+t7.7] w[+mC]=GMR57C10-GAL4}attP2 (BDSC # 39171) with males of either a GCaMP6m line w[1118]; P{y[+t7.7] w[+mC]=20XUAS-IVS-GCaMP6m}attP40 (BDSC # 42748) or an Arclight line w[*]; P{y[+t7.7] w[+mC]=UAS-ArcLight}attP40/CyO (BDSC # 51057). Adult F0 flies were allowed to lay eggs for 3-4 days and then larvae of the first, second, or third instar stage were separated from the food 1 to 4 days later. A 20% sucrose solution in water was added to the fly vial and floating larvae were collected after 10 minutes. Larval stage was identified based on size. For adult F1 flies, the larvae were allowed to mature to adulthood and adults were collected for a maximum of 7 days post-eclosion of the first F1 fly. Adults were then transferred weekly to new vials. Male and mated female adult flies of age 17 to 67 days were used in approximately equal proportions in the experiments.

### Reagents

A 20% sucrose solution in water was used to extract larvae from the food. Dissections and recordings were performed using physiological saline^55^ consisting of (in mM): 108 NaCl, 5 KCl, 4 MgCl2*6H2O, 3 CaCl2*2H2O, 1 NaH2PO4, 4 NaHCO3, 5 trehalose, 10 sucrose, 5 HEPES. pH was adjusted to 7.5 with NaOH. For some experiments, a 1.25 x KCl saline (6.25 mM) was used. All reagents were obtained from Sigma Aldrich (St. Louis, MO).

For determining a possible intrinsic temperature dependence of the GCaMP6m protein, brains from 16 adult flies less than 5 days old were dissected out in a calcium-free buffer solution consisting of (in mM): 10 HEPES, 2 EGTA, 135 NaCl. Brains were placed in 50 µL of the calcium-free buffer and homogenized using a homogenizer (Bel-Art ProCulture; Thermo Fisher Scientific, Waltham, MA). 1 µL of the homogenized brain solution was mixed with 1 µL of a 100 µM calcium buffer solution containing (in mM): 10 HEPES, 2 EGTA, 135 NaCl, 100 CaCl_2_*2H_2_O. Fluorescent imaging under 470 nm illumination light was performed on the final 50 µM brain solution as the temperature was cooled from 20 °C down to 0 °C. As a control to detect potential autofluorescence, the brain solution was imaged under 525 nm light at room temperature. No autofluorescence was observed. Finally, 2 µL of 50 µM calcium solution without brain tissue was imaged under 470 nm light. No fluorescence was observed.

### Dissection and Experiment Holders

Custom-made dissection dishes and animal holders were used during the experiments. For adults, a 3.75 x 2.5 x 0.025 cm (length x width x thickness) piece of aluminum was used as a base for securing the flies with dental cement (see Fig. 1). For larvae, 25 µm thick aluminum was used to create a 6.25 x 6.25 x 0.95 cm (length x width x height) square tray. A layer of 0.25 cm thick Sylgard 184 (Sigma, St. Louis, MO) was added to the holder. Dissections and mounting of animals were performed using a stereo microscope (Leica, Wetzlar Germany) with fluorescence adapter (Kramer Scientific, Amesbury, MA). Fluorescence excitation (470 nm) was provided by a Mightex BLS-Series High-Power LED Collimator Source (Mightex, Toronto, Ontario, Canada).

### Adult Cooling Experiments

Adult flies (1 - 4 animals) were secured to the adult fly holder using dental cement (Protemp, ESPE, St. Paul, MN). The wings and legs were placed in the cement to eliminate movement. A petroleum jelly well was created around the head and 0.5 mL of physiological saline was added. The abdomen and thorax of the fly remained outside the saline well. Care was taken to not obstruct the spiracles. The dorso-posterior head cuticle and trachea were removed to expose the brain. The holder with flies was then placed on the top of a 40 x 40 mm Peltier chip (Peltier Module TEC1-12706) that was controlled by a regulated power supply (Mastech HY3010Ex, Mastach. San Jose, CA). A water-cooling block below the Peltier chip ensured efficient heat transfer (see Fig. 1). A temperature logger (TC0520, PerfectPrime, New York, NY) recording at 0.2 Hz was placed into the physiological saline well.

### Larval Cooling Experiments

For high-throughput experiments, up to 16 larvae were secured to the larval fly holder. Each larva was secured using 2 pins, one near the mouth hooks and another near the spiracles at the posterior end. A 0.2 mL drop of physiological saline was added to cover all larvae on the holder. All animals on the holder were imaged simultaneously (see Fig. 6A).

For fillet preparations, larvae were pinned with their dorsal side up using 2 pins, one near the mouth hooks and another near the spiracles at the posterior end (Fig. 6B). A 0.2 mL drop of physiological saline was added to cover all larvae on the holder. Two small transversal cuts were made at the anterior and posterior ends of each larva, and a longitudinal incision was made along the dorsal body wall of the animal to fillet the animal. Additional pins were used to open and secure the body wall. The gut and trachea were removed to expose the brain. As in the case with the adults, the holder with the larvae was placed on a water-cooled Peltier chip. A temperature probe recording at 0.2 Hz was placed into the physiological saline surrounding the larvae. The temperature was adjusted by changing the voltage supplied to the Peltier chip.

### Potassium Chloride Application Experiments

To test whether extracellular KCl application could elicit SD, we increased the potassium concentration in the saline bath that surrounded the fly brain (in adults) or the brain hemispheres and ventral nerve cord (in larvae). A drop (4µl) of 1M KCl solution was added to the bath, and the location and time of application were noted. Fluorescence imaging continued throughout the application. The application rapidly increased the potassium concentration from the normal 5 mM to 12.9 mM.

### Electrophysiology

Transperineuronal potentials were obtained by impaling the adult brain using 20-30 MΩ glass microelectrodes filled with 0.6 M K_2_SO_4_ + 20 mM KCl electrolyte solution. Electrodes were pulled using a Sutter P97 puller (Sutter Instruments, Novato, CA). Signals were filtered and amplified through an Axoclamp 900A amplifier (Molecular Devices, San Jose, CA) in bridge mode. Files were recorded, saved, and analyzed using Spike2 Software at 10 kHz (version 7.18; CED, Cambridge, UK) and a Power 1401 (CED).

### Fluorescent Imaging and Analysis

Fluorescence data were recorded at 10 Hz using an Olympus-BX51 epifluorescence microscope with either a UMPLFLN 10XW (high-resolution experiments; 0.3NA) or MPLFLN 4X objective (high-throughput experiments; 0.15NA), CoolLed PE-4000 fluorescence illuminator (470nm; 505nm excitation cut-off filter, 525/50nm emission filter), and Basler ace acA4024-29um camera (Basler, Ahrensburg, Germany). Images were imported into FIJI^56^ and regions of interest (ROIs) were drawn around either the entire brain (to determine start and end of SD) or small regions of the brain (for propagation analysis). The mean fluorescence value for each ROI was plotted as a function of time, which was then correlated with temperature. A custom-made script was used to identify the starting and ending times of SD. The ending time was defined to be the time of maximum fluorescence. The starting time was defined to be the time where fluorescence began to increase rapidly. This point was found by shifting and rotating the coordinate axes of the fluorescence vs. time curve (see supplemental 7). A new origin was defined at the initial fluorescence value (t = 0) and the x- and y-axes were rotated such that the new y-axis was along a line from the new origin to the maximum of the original fluorescence vs. time curve. The sharp change in slope identifying the start of SD could then be found from the point in the fluorescence curve that had the largest x-value in the new coordinate system.

### Statistical Analysis and Figure Production

For normally distributed and paired data, either paired t-tests (for comparison of two conditions) or repeated measures ANOVA with Student-Neuman-Keuls (SNK) posthoc tests at P<0.05. For other paired data, nonparametric repeated measures ANOVA on Ranks tests were used. For unpaired normally distributed data, either t-tests (for two group comparisons) or One-Way ANOVAs with SNK posthoc tests were use. For other unpaired data, One-Way ANOVAs on Ranks were used. Statistical tests were calculated with Sigmaplot (v15, Grafiti LLC, Palo Alto, CA). Unless stated otherwise, data are presented as mean ± SEM. Alternatively, individual data points for each animal are given. Significant differences are stated as *p<0.05, **p<0.01, ***p<0.001.

Figures were prepared with CorelDraw X7 (Corel Cooperation, Ottawa, Canada), Excel 365 (Microsoft, Redmond, WA), Adobe (San Jose, CA), SigmaPlot (version 11, Systat Software, San Jose, CA), and VSDC video editor (Flash-Integro LLC, Tashkent, Uzbekistan).

## Supporting information

Supplemental figures and description

## V. Acknowledgements.

We would like to thank the College of Arts and Sciences and the Physics Department at Illinois State University for providing financial support. We would also like to thank the School of Biological Sciences for providing laboratory space and commodities. Further financial support through Firebird and Birdfeeder grants was provided by the Office of Student Research at Illinois State University. Further thanks go to Pedro Galvan and Abigail Spena for carrying out preliminary experiments. Stocks obtained from the Bloomington Drosophila Stock Center (NIH P40OD018537) were used in this study.

