## Supplemental figures and description for "Whole brain fluorescence imaging in *Drosophila* reveals spreading depression and its initiation, propagation, and resilience dynamics"

*Supplemental 1*: False-color video of spontaneous activity in the adult fly brain. GCaMP6m was expressed pan-neuronally. Warmer colors represent higher fluorescence. The video is shown at five times the original speed. Anterior is to the top.

*Supplemental 2*: False-color video of cooling-induced spreading wave of fluorescence in the adult fly brain. GCaMP6m was expressed pan-neuronally. Warmer colors represent higher fluorescence. The video is shown at five times the original speed. Anterior is to the top.

*Supplemental 3.* Changes in GCaMP6m fluorescence during cooling and warming of intact brains (darker colors) and homogenized brains (lighter color). No rapid or large changes in fluorescence were detected in homogenized brains.

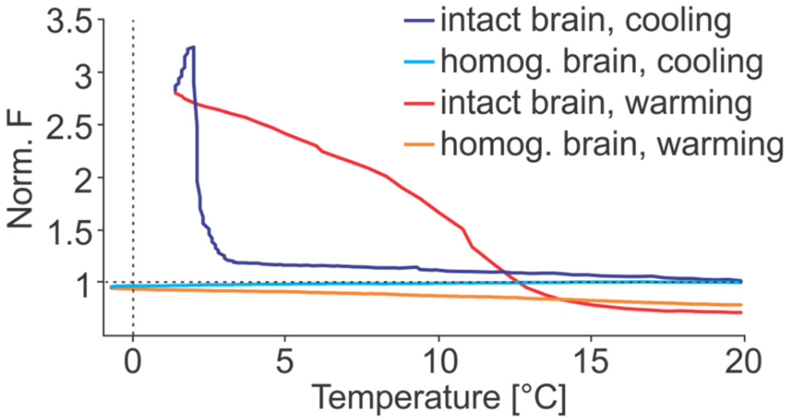


*Supplemental 4*: False-color video of spontaneous activity in the third larval instar ventral nerve cord. GCaMP6m was expressed pan-neuronally. Warmer colors represent higher fluorescence. The video is shown at five times the original speed. Anterior is to the bottom left.

*Supplemental 5*: False-color video of cooling-induced spreading wave of fluorescence in the third larval instar ventral nerve cord. GCaMP6m was expressed pan-neuronally. Warmer colors represent higher fluorescence. The video is shown at five times the original speed. Anterior is to the top left.

*Supplemental 6.* SD confers short-term resilience to subsequent SDs across developmental stages of the fly nervous system (for detailed statistics, see text). A) Quantification of SD temperature of multiple SDs that were elicited by repeated cooling of first instar nervous systems. Subsequent SDs required colder temperatures to elicit SD. B) Quantification for third instar larvae. C) Quantification for adult brains.

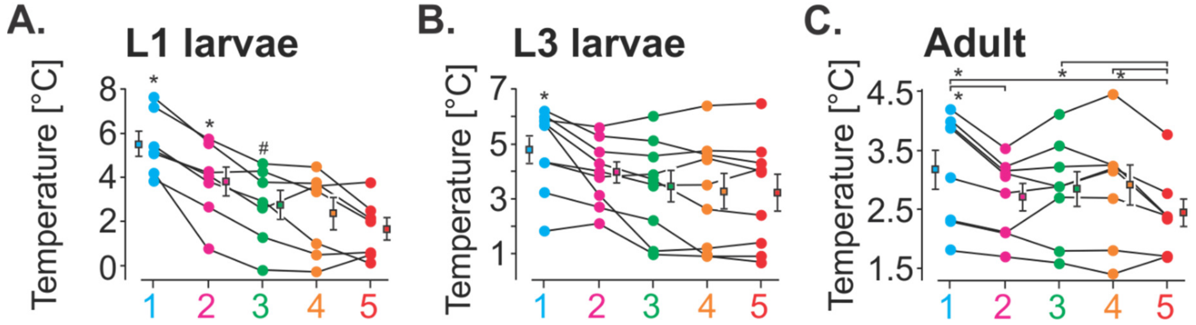

*Supplemental 7.* Schematic illustration of method used to find SD start time. The black line shows the total fluorescence over time with the original coordinate axes in black. A new coordinate system (red) was defined with its origin at the initial fluorescence value and the y-axis along a line from the new origin to the maximum of the fluorescence curve. The point of maximum transverse distance of the fluorescence curve from the new y-axis (blue arrow) identifies the start of SD. The end of SD was identified as the time of maximum fluorescence (purple arrow).

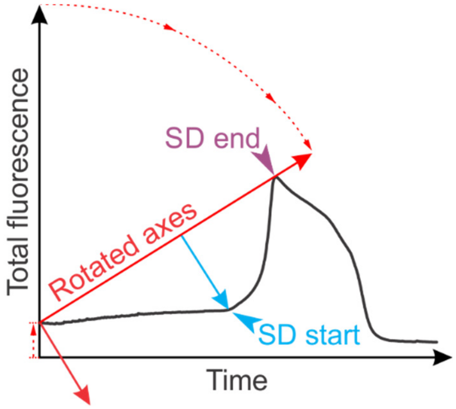
